# Anthrax intoxication reveals that ER-Golgi membrane contact sites control the formation of plasma membrane lipid nanodomains

**DOI:** 10.1101/2022.05.18.492252

**Authors:** Muhammad U. Anwar, Oksana A. Sergeeva, Laurence Abrami, Francisco Mesquita, Ilya Lukonin, Triana Amen, Audrey Chuat, Laura Capolupo, Prisca Liberali, Giovanni D’Angelo, F. Gisou van der Goot

## Abstract

To promote infections, pathogens exploit host cell machineries including structural elements of the plasma membrane. Studying these interactions and identifying involved molecular players is an ideal way to gain insights into the fundamental biology of the host cell. Here, using the anthrax toxin, we screened a 1500-gene library of regulatory, cell surface, and membrane trafficking genes for their involvement in the intoxication process. We found that the ER–Golgi-localized proteins TMED2 and TMED10 are required for toxin oligomerization at the cell surface, an essential step for anthrax intoxication that depends on localization to cholesterol-rich lipid nanodomains. Further biochemical, morphological and mechanistic analyses showed that TMED2 and TMED10 are essential components of a multiprotein supercomplex that operates exchange of both cholesterol and ceramides at ER-Golgi membrane contact sites. Overall, this study of anthrax intoxication led to the discovery that lipid compositional remodelling at ER-Golgi interfaces fully controls the formation of functional membrane nanodomains at the cell surface.

## Main

Pathogens have evolved to co-opt existing cellular processes of their hosts. Consequently, studies of host-pathogen interactions are continuously deepening our understanding of fundamental biological processes and have helped uncover ones such as the fusion of synaptic vesicles, dynamics of the actin cytoskeleton, or retrograde transport from the Golgi to the ER (Mañes et al., 2003; Mesquita et al., 2020; Schiavo & van der Goot, 2001). Here, we searched for genes enabling anthrax intoxication, which uncovered proteins involved in compartmentalisation of the plasma membrane.

Anthrax toxin is an exotoxin secreted by virulent strains of *Bacillus anthracis* and is composed of three polypeptides, two enzymatic subunits – lethal factor (LF) and edema factor (EF) – and one receptor binding subunit, protective antigen (PA). During the intoxication process (Figure 1A), the monomeric 83 kDa precursor of PA (PA^83^) first binds to one of two toxin receptors, CMG2 and TEM8, at the cell surface (Friebe et al., 2016). PA^83^ is then cleaved via proprotein convertases, such as furin (Klimpel et al., 1992; Sergeeva & van der Goot, 2019). This process is enhanced when the receptor-bound PA and the protease are brought together through association with cholesterol-rich nanodomains (Abrami et al., 2003; Levental et al., 2020).

**Figure 1:**
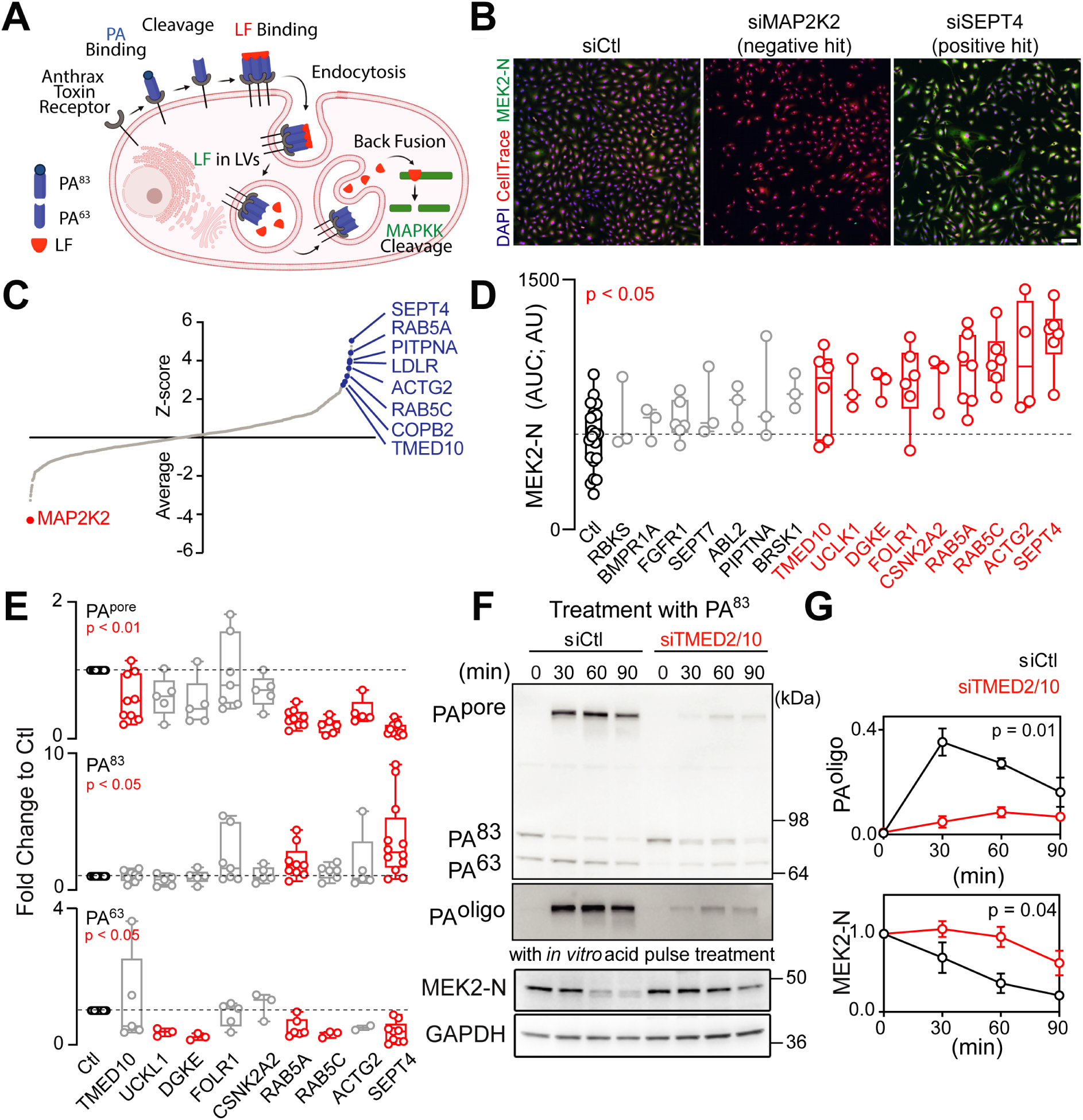
TMED2/10 are required for anthrax intoxication: See also Figure S1. **A)** Schematic representation of anthrax intoxication. **B)** Representative IF images obtained in the image-based screen for anthrax intoxication. siSEPT4 and siMAP2K2 are examples of positive and negative hits, respectively. CellTrace, red; anti MEK2-N antibody, green; Scale bar is 100 µm. **C)** Image-based screen results sorted by increasing Z-score. Representative genes whose knockdown impaired MEK2 cleavage are in blue; MAP2K2 (a.k.a. MEK2) knockdown served as a specificity control (in red). **D)** Effect of the knockdown of selected hits from the screening on MEK2 cleavage as assessed by western blot in a 0 to 90 min intoxication time course, with each point representing the area under the curve for normalized MEK2 values. Red represents conditions where MEK2 cleavage was significantly impaired (p < 0.05). **E)** Effect of silencing hit genes validated in (D) on the levels of PA^63^ (lower panel), PA^83^ (middle panel), and PA^pore^ (upper panel). Red represents conditions associated with significant changes. **F).** Western blot of control and siTMED2/10-treated cells after anthrax toxin treatment for the indicated times. Membranes were probed with antibodies against PA, MEK2-N, and GAPDH. **G)** Quantification of PA^oligo^ rendered SDS-resistant by acidification of the cell lysate (upper panel) and MEK2-N cleavage (lower panel) from the western blot for both conditions.

The cleaved 63 kDa PA (PA^63^) oligomerizes into a ring-like structure (PA^oligo^) (Milne et al., 1994) that binds the enzymatic subunits, LF and EF (Friebe et al., 2016). The hetero-oligomeric toxin-receptor complex is then internalized and trafficked to late endosomes (Abrami, Bischofberger, et al., 2010; Abrami et al., 2004, 2006; Abrami, Kunz, et al., 2010) (Figure 1A), where the acidic environment triggers membrane insertion of the PA oligomer. This forms a membrane translocation pore (PA^pore^) through which EF and LF are transported (Krantz et al., 2006; Sun & Jacquez, 2016) ultimately allowing them access to the cytosol where they exert their toxic roles: LF cleaving MAP kinase kinases (MAPKK) and EF increasing cAMP levels (Friebe et al., 2016).

While the global view of the mechanism of anthrax intoxication is available, the molecular players of many steps are missing. We therefore undertook an image-based RNA interference (RNAi) screen to identify novel modulators of the toxin uptake route. Several factors affected PA cleavage and pore formation, indicating that these early steps require the action of multiple cellular proteins. Unexpectedly, downregulating two proteins in the early secretory pathway, TMED2 and TMED10, prevented PA oligomerization at the plasma membrane. Intriguingly,TMED2/10 silencing led to loss of surface cholesterol-rich nanodomains (Levental et al., 2020) required for the toxin’s mode of action. In depth analysis showed that these two proteins act as essential organizers of large protein supercomplexes at ER-Golgi membrane contact sites, responsible for the specific transfer of cholesterol and ceramide between the two organelles. Thus, by screening for molecular players in the anthrax intoxication process, we retrieved key components of the molecular machinery necessary for the formation of functional plasma membrane lipid domains relevant for pathogen infection, and for fundamental physiological processes.

## Results and discussion

### TMED2/10 are required for anthrax intoxication

To search for genes involved in anthrax intoxication, we developed an image-based screen that employs the N-terminal cleavage of the MAPKK MEK2 as a readout, as MAPKKs are cleaved by LF upon productive cell intoxication (Park et al., 2002) (Figure 1A). For this screen, we chose an siRNA library against 1500 genes, which included a validated library of endocytosis and kinase genes with additional siRNAs against cell-surface proteins (Liberali et al., 2014). Cells were first silenced with siRNAs, incubated with the two anthrax toxin components (PA and LF), fixed, and stained using an antibody against the N-terminus of MEK2 (MEK2-N). The MEK2-N signal decreased in control cells, while remaining high when silencing genes affecting anthrax toxin entry (Figure 1B). We identified 94 silencing conditions with significantly higher MEK2-N levels than in controls (Z-score > 2; Figure 1C).

Hypothesising that toxin entry relied on the structural integrity of the endocytic organelles, we considered in parallel the morphology of the early endosomal compartment (i.e., assessing the distribution of the early endosomal marker EEA1; Figure S1A) that uncovered 85 silencing conditions with significantly altered endosomal morphology (Z-score > 2 or < -2; Table S1). Interestingly, hits from the EEA1 and toxin entry screens differed largely, with only 16 silencing conditions impacting both endosomal morphology and anthrax intoxication (randomly expected common hits 5.7). This suggests that the molecular machinery assisting anthrax toxin endocytosis is distinct from that responsible for maintaining endosomal morphology (Figure 1B).

We next compared the results of the anthrax toxin entry screen to those of the endocytome screens, which tested the dependence of multiple endocytic pathways on signalling and trafficking genes (Liberali et al., 2014). The endocytome screens yielded 5 categories of endocytic processes (i.e., cholera toxin [ChTxB] uptake, transferrin endocytosis, fluid phase endocytosis, EGF endocytosis, and micropinocytosis/LDL endocytosis) that depended on discrete regulatory gene modules (Liberali et al., 2014). We mapped the Z-scores associated with the anthrax toxin entry screen on these modules and observed that silencing the genes regulating EGF endocytosis reduced anthrax toxin-induced MEK2 cleavage (Figure S1C). This suggests that anthrax toxin entry has general molecular requirements similar to those of EGF endocytosis, consistent with previous findings (Abrami et al., 2003, 2004, 2006).

Based on high Z-scores, detectable expression in RPE1 cells, and efficient silencing by siRNA, we chose the 14 best hits (Table S2). Anthrax toxin entry can be followed by monitoring (Figure 1A): 1. Binding of PA^83^ and its subsequent cleavage into PA^63^, which occur at the cell surface; 2. Formation of the SDS-resistant oligomeric PA^pore^, which occurs in the acidic endosomes; and 3. MEK2 cleavage, revealing the release of LF into the cytosol. All three steps were monitored using western blotting against PA or MEK2-N, either at a single time point (30 min) or as a function of time.

Silencing the candidates had no effect on the surface expression of the toxin receptor CMG2 (Figure S1D, E), whereas silencing 9 of these reduced the kinetics of MEK2 cleavage (Figure 1D). Silencing 5 candidates (ACTG2, RAB5A, RAB5C, SEPT4, and TMED10) inhibited the formation of PA^pore,^ and silencing 2 (SEPT4 and RAB5A) increased the amount of cell-bound PA^83^ due to diminished cleavage into PA^63^ (Figure 1E). TMED10 stood out as particularly interesting because it is reported as a protein of the early secretory pathway, yet its silencing inhibited the formation of PA^pore^ without affecting the amount of anthrax toxin receptor at the plasma membrane, nor toxin cleavage (Figure 1E and Figure S1D, E).

TMED10 belongs to a family of type I transmembrane proteins with a large luminal domain and a short cytosolic tail (Emery et al., 1999; D. Gommel et al., 1999). TMED members can hetero-oligomerize, and TMED10 in particular has been shown to dimerize with TMED2 (Zavodszky & Hegde, 2019). Consistently, silencing TMED2 or TMED2 and TMED10 together inhibited LF-dependent MEK2 cleavage and the formation of PA^pore^ in endosomes (Figure 1F, G and Figure S1F, H). For all further studies, we therefore silenced both TMED 2 and 10 simultaneously.

In addressing the role of TMED2 and 10 in pore formation, we found that this is due to the failure of PA oligomerization in TMED2 and/ or 10-depleted cells (Figure 1F, G and Figure S1F, G). This was monitored an *in vitro* acid pulse, which treats cell lysates with an acidic buffer, to trigger conversion of all oligomers to the SDS-resistant PA^pore^. To further confirm that TMED2/10 affect PA oligomerization, we converted PA^83^ to PA^63^ *in vitro* using trypsin prior to its addition to cells. PA^pore^ formation was still delayed (Figure S1I). Overall, these observations indicate that TMED2/10 specifically affect PA oligomerization at the cell surface. This is the first time that a protein was found to affect this specific step of anthrax intoxication.

### TMED2/10 affect the formation of functional plasma membrane lipid nanodomains

As PA^63^ oligomerization requires association with cholesterol-rich lipid nanodomains (Abrami et al., 2003; Sergeeva & van der Goot, 2019), we next investigated this association by exploiting nanodomain resistance to solubilization at 4°C in specific detergents, such as Triton X-100, hence their name of detergent-resistant membranes (DRMs) (Levental et al., 2020). Strikingly, while PA^63^ was retrieved in the DRM fraction (fraction 2) of control cells as previously shown, it was solely found in the detergent-soluble fractions (5 and 6) of TMED2/10-silenced cells. This was not due to an off-target effect of the siRNA, since it could be reverted by the recomplementation with siRNA-resistant TMED2 and/or TMED10 constructs (Figure 2A, B).

**Figure 2.**
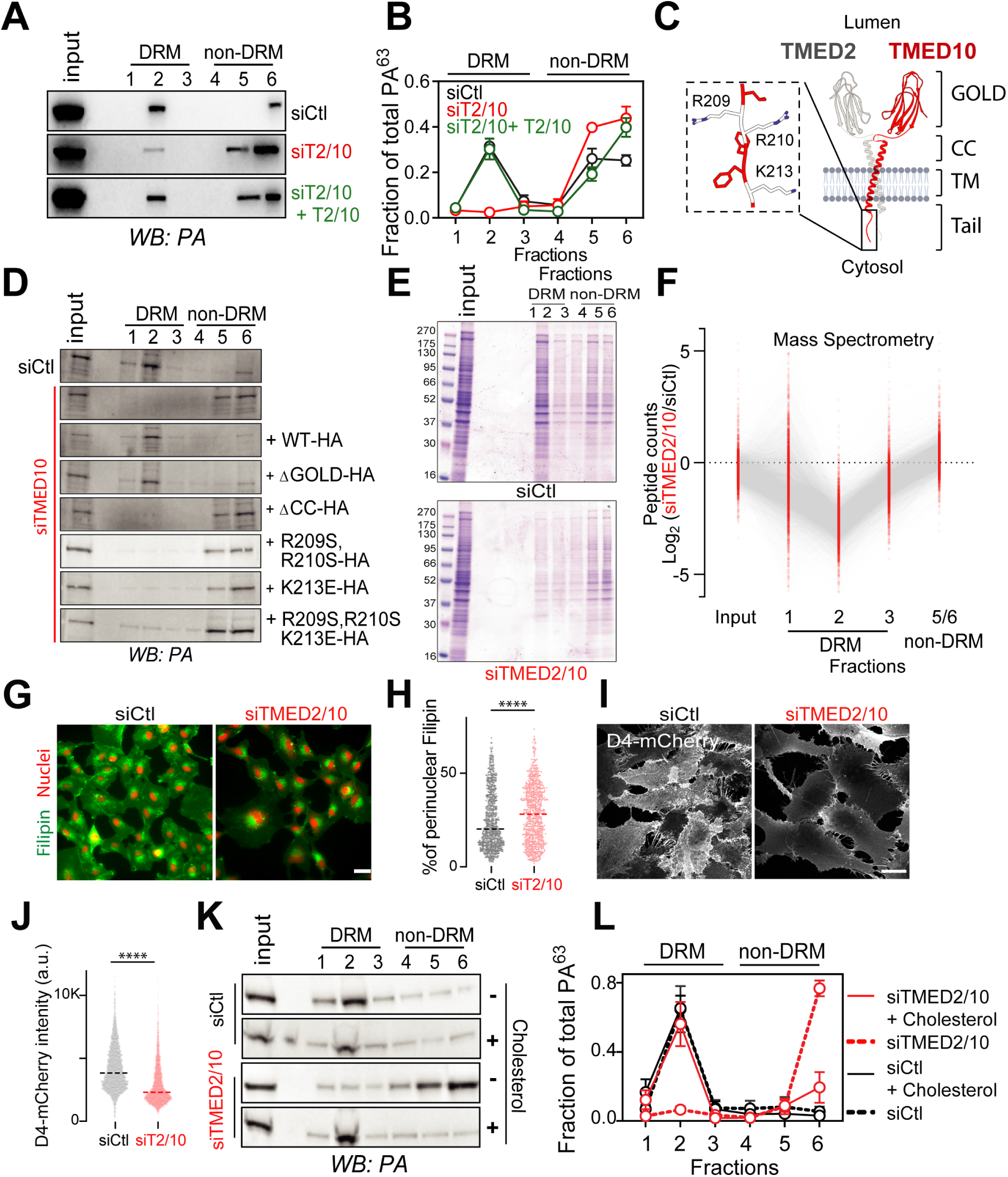
TMEDs are required for the assembly of lipid nanodomains. See also Figure S2. **A)** Western blot of step gradient fractions in control and siTMED2/10 cells with or without TMED2/10 overexpression. Silenced or Silenced + TMED2/10 transfected cells were lysed with Triton-X. The lysates were then fractionated using an Optiprep step gradient, and an equal volume from each fraction was loaded on the gel (see methods). Membranes were probed with anti-PA. **B)** Values from the western blot in (A) were normalised within each experiment. Detergent-resistant membranes (DRM) or non-DRM fractions are indicated above the lanes. **C)** Schematic representation of TMED2/10 hetero-dimer with main domains: GOLgi Dynamics (GOLD), coiled coil (CC), transmembrane (TM), and cytoplasmic tail. Positively charged residues are labelled within the tail (magnified view). The lumen could either be Golgi or ER. **D)** Same as (A) with overexpression of different TMED10 mutants (see also Figure S2A). **E)** Same as in (A), fractions were run on SDS-PAGE and gels were stained with Coomassie. **F)** Same as in (A), fractions were submitted for mass spec either as fraction 1, 2, 3, or an equal combination of 5 and 6 (labelled 5/6). Each point represents the log2 ratio of one protein’s abundance in siTMED2/10 over siCtl. **G)** Silenced cells were fixed and stained with filipin and NucGreen and were imaged. **H)** Each cell from (G) was segmented and quantified for its total and perinuclear average integrated filipin signal intensity. The median of the proportion of the perinuclear to total filipin signal was graphed. n = 881, siCtl; n = 772, siTMED2/10 from three replicates. **I)** Control and silenced TMED2/10 cells were fixed and stained with D4-mCherry and imaged using laser scanning confocal microscopy. **J)** Average intensity from cells in (I) of surface mCherry is plotted per cell per condition. n = 5078, siCtl; n = 6151, siTMED2/10 from three replicates. **K)** and **L)** Same as in (A) and (B), the silenced cells were either untreated or nourished with cholesterol through βCD-Cholesterol (1 mM) treatment for 24 h. Results are mean ± SEM (n = 3). For (g-k) *p < 0.05, ****p < 0.0001 (Unpaired two-tailed t-test). All scale bars are 10 µm.

We next examined which domain of TMED10 was responsible for the recovery of PA in the DRM fraction by recomplementing siRNA-silenced cells with modified TMED10 constructs lacking specific domains (Figure 2C). Recomplementation with a construct lacking the N-terminal luminal GOLD domain, involved in the binding to glycosylphosphatidylinositol-anchored proteins allowed recovery of the DRM- associated anthrax PA. In contrast, when expressing constructs missing the coiled-coil domain, involved in dimerization of TMEDs (Zavodszky & Hegde, 2019), PA remained detergent soluble (Figure 2D and Figure S2A). Mutants with modifications, even single point mutations, to the charged residues of the cytosolic tail, also did not permit recovery of DRM-associated PA (Figure 2D and Figure S2A). Thus, the abilities of TMED10 to dimerize as well as to organise interactions via its cytosolic tail (D. U. Gommel, 2001; Zavodszky & Hegde, 2019) are necessary to recover PA in the DRM fraction.

A decreased presence in fraction 2 of the gradient in TMED2/10-silenced cells was not specific to PA; the levels of other proteins commonly found in nanodomains, such as flotillin and CMG2, also decreased significantly (Figure S2B). Global assessment of protein association to the different fractions of the gradient by Coomassie staining and mass spectrometry analysis (Figure 2E, F) confirmed a generalised and profound reduction in fraction 2 in TMED2/10-silenced cells. These observations suggest that TMED2/10 play a widespread role in establishing lipid domains rather than specifically affecting PA association with them.

Cholesterol is an indispensable component of nanodomains (Levental et al., 2020), and we have previously shown that extracting cholesterol with methyl-ß-cyclodextrin (mßCD) affected anthrax toxin action (Abrami et al., 2003). We thus investigated the effects of TMED2/10-silencing on the cellular distribution of cholesterol. We visualised cholesterol by labelling cells with the fluorescent cholesterol-binding fungal metabolite Filipin III (Maxfield & Wüstner, 2012; Whitfield et al., 1955). The labelling was strikingly different in TMED2/10-silenced cells as compared to controls: instead of a marked plasma membrane staining, cholesterol accumulated in the perinuclear region (Figure 2G, H). The loss of cholesterol from the plasma membrane was confirmed by staining non-permeabilized cells with a probe that binds accessible cholesterol, D4-mCherry (Ramachandran et al., 2002) (see methods for details) (Figure 2I, J).

As TMED2 has been reported to bind sphingomyelin (Contreras et al., 2012), also a component of nanodomains (Levental et al., 2020), and to affect sphingomyelin levels (Jiménez-Rojo et al., 2020), we tested whether the prsence of sphingomyelin was reduced at the cell surface of TMED2/10-silenced cells. Labelling non-permeabilized cells with the earthworm sphingomyelin-binding toxin Lysenin, however, indicated that, in our conditions, surface sphingomyelin levels were not substantially different between control and silenced cells (Figure 2C).

We next tested whether the reduction in cholesterol at the plasma membrane is the main reason for the observed phenotype. Exogenous cholesterol was added by treating cells with 1 mM of methyl-ß-cyclodextrin/cholesterol complexes (mßCD - Chol). This treatment was sufficient to recover PA in fraction 2 (Figure 2L, M), indicating that the reduction of surface cholesterol is the major reason for the loss of DRM-associated PA in TMED2/10-silenced cells.

### TMED2/10 affect lipid fluxes through the secretory pathway

Because cholesterol levels at the plasma membrane are altered upon TMED2/10 silencing, we wondered if cholesterol metabolism is compromised in TMED2/10-KD cells. Lipidomics experiments showed that cholesteryl esters were substantially elevated upon TMED2/10 silencing, while no significant changes were observed in overall cholesterol levels (Figure 3A, B). Accordingly, the size and number of lipid droplets (where cholesteryl esters are stored) increased upon TMED2/10 silencing (Figure 3C, D).

**Figure 3.**
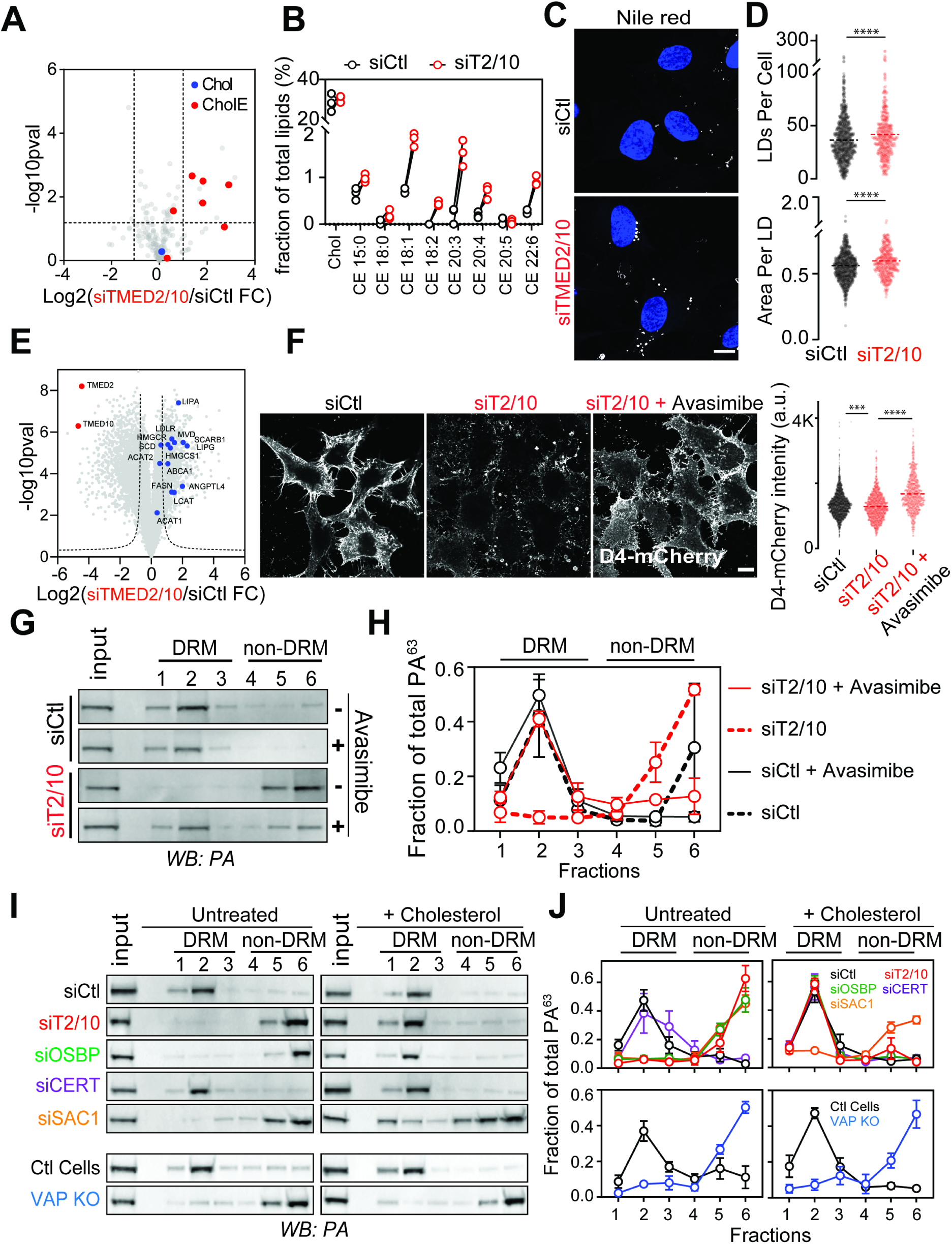
TMED2/10 affect cholesterol transport to the cell surface. See also Figure S3. **A)** Lipidomic analysis of siTMED2/10 *vs*. siCtl cells. The cholesterol and cholesteryl esters (CE) *spp.* are highlighted in terms of log2 of the siTMED2/10 over siCtl ratio. **B)** Levels of Chol and different CE *spp.* In siCtl and siT2/10. **C)** Control and silenced TMED2/10 cells were fixed and stained with Hoechst (nuclei, blue) and Nile Red (LDs, gray) and imaged using laser scanning confocal microscopy. **D)** LDs in each condition were segmented based on the Nile Red signal. Average intensity and droplet size were calculated per cell in each condition. n = 1140, siCtl; n = 449, siTMED2/10 from three replicates. **E)** RNA-Seq analysis of siTMED2/10 *vs*. siCtl cells. Genes involved in cholesterol transport and synthesis are highlighted in blue as log2 of the siTMED2/10 *vs.* siCtl ratio. **F)** HeLa control and TMED2/10-silenced cells were fixed and stained for D4-mCherry. Cells were either treated with Avasimibe (10 uM) or not (Ctl). Average intensity of surface mCherry from the images is plotted per cell per condition. n = 1676, siCtl; n = 1102, siTMED2/10; n = 786, siTMED2/10 + Avasimibe from three replicates. **G)** Western blot of step-gradient fractions in control and siTMED2/10 cells with or without ACAT inhibition. Cells were lysed with Triton-X and placed below an Optiprep step gradient and ultra-centrifuged for 2 h. Equal volume fractions were taken from the top (1) to the bottom (6) and loaded on gels. Membranes were probed with anti-PA. **H)** The values from (G) were normalized within each experiment. DRM or non-DRM fractions are indicated above each lane. **I)** and **J).** Same as in (G) and (H), silenced cells were either treated with mßCD-Chol (1 mM) or not (untreated). All scale bars are 10 µm.

RNASeq analysis of TMED2/10-silenced cells consistently showed that acylCoA:cholesterol acyltranferases (ACATs) 1 and 2 were upregulated. In fact, various genes involved in cholesterol biosynthesis or uptake were increased, such as HMG-CoA synthase 1 (HMGCS1), HMG-CoA reductase (HMGCR), diphosphomevalonate decarboxylase (MVD), and stearoyl-CoA desaturase (SCD1) (Figure 3E). The increase could also be observed at the protein level (Figure S3A, B). The LDL receptor gene was also upregulated and its subcellular localization remained unchanged in silenced cells (Figure S3C, D), indicating that defective cholesterol uptake is unlikely to play a role in the observed phenotypes.

As TMED2/10 silencing impacts sterol esterification, we tested whether inhibiting ACATs might restore the flux of cholesterol towards the plasma membrane. Upon ACAT inhibition, we indeed observed a recovery of plasma membrane levels of cholesterol as assessed by D4-mCherry (Figure 3F) and the concomitant DRM-association of PA (Figure 3G, H).

We conclude that silencing TMED2/10 impairs cholesterol transport to the plasma membrane and promotes its storage in lipid droplets. These effects prevent the formation of plasma membrane lipid nanodomains and thus the oligomerization of PA.

### Membrane contact site components are involved in plasma membrane nanodomain formation

Cholesterol is synthesised in the ER, and is then transported to the Golgi, either in vesicles or, more substantially, via the oxysterol binding protein (OSBP), a lipid-transfer protein that localizes at membrane contact sites (MCSs) between the two compartments (Mesmin et al., 2013, 2019). Cholesterol is subsequently trafficked to the plasma membrane and the endocytic pathway by vesicular carriers (Antonny et al., 2018; Luo et al., 2020) Upon OSBP inhibition, cholesterol transport to the Golgi is diminished, undergoes esterification in the ER and is stored in lipid droplets (Luo et al., 2020). OSBP also ensures the transport of phosphatidylinositol-4-phosphate [PtdIns(4)*P*] from the Golgi to the ER for its dephosphorylation, serving as a counterflow for cholesterol transfer (Antonny et al., 2018; Hanada et al., 2009). Interestingly, in TMED2/10 silenced cells, this phosphoinositide accumulated on the Golgi (Figure S3E, F) phenocopying OSBP inhibition.

Finally, sites of contact between ER and Golgi also mediate transfer of ceramide for its conversion to sphingomyelin. This is operated by the cytosolic ceramide transporter (CERT) (Hanada et al., 2009). Upon TMED2/10 silencing, ceramide conversion to sphingomyelin was impaired and glucosylceramide synthesis and steady-state levels were strongly increased (Figure S3G, H), phenocopying CERT inhibition (Capasso et al., 2017; Hanada et al., 2003, 2009).

The high level of esterified cholesterol, the Golgi accumulation of PtdIns(4)*P* and the reduced sphingomyelin production in TMED2/10-silenced cells raised the hypothesis that these proteins play a key role in lipid exchange at MCSs between the ER and the Golgi, two compartments between which they are known to cycle (Strating & Martens, 2009). This hypothesis predicts that established MCS components, when silenced, should similarly affect the recovery of PA in DRM fractions as silencing TMED2/10. OSBP association with the ER is mediated by the dimeric transmembrane protein VAPA, while its association with the Golgi occurs through ARF1 and PtdIns(4)*P* (Antonny et al., 2018; Levine & Munro, 2002; Mesmin et al., 2013). The PtdIns(4)*P* phosphatase SAC1, which ensures that the phosphoinositide is consumed when transported to the ER (Mesmin et al., 2019) is also a key player of MCSs, since PtdIns(4)*P* has been proposed to be the energy source in the transport of cholesterol against its concentration gradient. Silencing SAC1 and OSBP, but not CERT led to a loss of DRM-associated PA. The phenotype of OSBP silencing could again be reverted by the exogenous addition of cholesterol (Figure 3I, J).

To study the role of VAPs, we used an available double knockout cell line for VAPA and VAPB (Dong et al., 2016). Here, PA was lost from fraction 2, and this association could only be moderately recovered by cholesterol addition (Figure 3I, J). Consistently, D4 staining (Figure S3I, J) showed that surface cholesterol levels were significantly reduced in VAPA/B knockout cells. Differently from TMED2/10-KD cells, VAPA/B knockout cells showed a substantial decrease in plasma membrane sphingomyelin as assessed by Lysenin staining (Figure S3K). The lack of DRM recovery suggests that cholesterol, in the absence of sphingolipids, is insufficient for the formation of surface nanodomains. VAPA/B knockout cells also differed from TMED2/10 silenced cells in that ACATs and SCD1 were not upregulated (Figure S3L, M).

Thus, MCS components involved in cholesterol transfer between the ER and the Golgi, i.e. VAPA/B, OSBP and SAC1, ensure the presence of cholesterol at the cell surface and the formation of nanodomains to which PA can associate.

### TMED2/10 silencing does not affect the localisation of ER-Golgi MCS components

Since TMED proteins have been shown to be cargo protein receptors, a possibility is that they are required for MCS components to properly localize. We first looked at the localization of TMED2 and 10 in RPE1 cells, where co-localization with Golgi markers such as Giantin (medial-Golgi), GM130 (cis-Golgi) (Figure 4A), and GOLPH3 (trans-Golgi) indicate accumulation in the Golgi (Figure S4A). Similar localization was observed in HeLa cells (Figure S4A). Co-localization with Golgi markers was previously reported (Mesmin et al., 2013, 2019) for the MCS proteins OSBP, CERT, and the CERT phosphatase PPM1L, which dephosphorylates CERT to enhance its membrane association (Saito et al., 2008) while VAPA and SAC1 are well established to localise to the ER, all consistent with our observations (Figure 4A). Silencing TMED2/10 did not grossly affect the localization of these MCS components (Figure 4A). The staining did appear different, however, due to an alteration of the Golgi morphology (Figure 4B, C), which was consistently fragmented.

**Figure 4.**
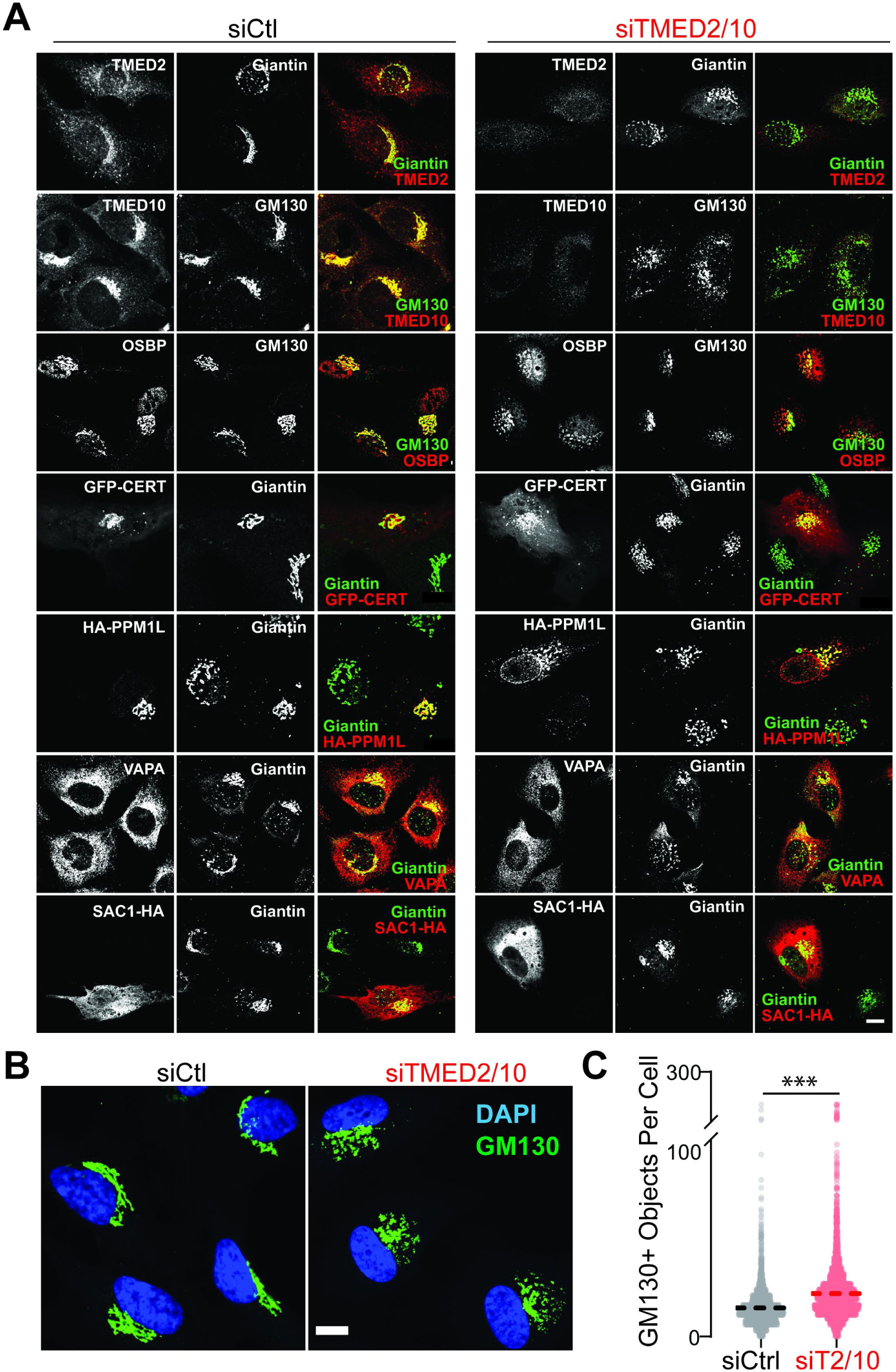
TMED2/10 affect Golgi morphology. See also Figure S4. **A)** Control or silenced cells were fixed and permeabilized 72 h post siRNA transfections and stained for different MCS proteins and Golgi markers. Tagged versions of PPM1L, CERT, and SAC1 were overexpressed 24 h pre-fixation in all cells. Images were acquired using laser confocal microscopy. **B)** Same as in (A). **C)** Cells were segmented using their labelled nuclei and their GM130-positive objects were quantified. n = 4327, siCtl; n = 8369, siTMED2/10 from three replicates. The average Golgi fragments per cell are shown. An unpaired two-tailed t-test was conducted to calculate the significance. Results are mean ± SEM. All scale bars are 10 µm.

Fragmentation of the Golgi has been previously reported upon upregulation of the stearoyl-CoA desaturase SCD1 (Lita et al., 2021). Since SCD1 was upregulated in TMED2/10 silenced cells, we tested the effect of SCD1 inhibition (A939572) on the Golgi morphology in HeLa cells, which partially reverted the phenotype (Figure S4B, C). Since our previous data showed a recovery of plasma membrane cholesterol levels upon ACAT inhibition (Figure 3E, F), we asked whether this inhibition would also restore the Golgi morphology. Treatment of silenced cells with Avasimibe indeed showed a significant reversion of Golgi fragmentation (Figure S4B, C).

Overall, our biochemical (Figure 2–3 and Figure S2–3) and morphological analyses (Figure 4 and Figure S4) suggest that TMED2/10 are directly involved in the transport of cholesterol at ER-Golgi MCSs.

### TMED2/10 are required for the assembly of ER-Golgi MCS protein supercomplexes

Finally, we analysed whether TMED2/10 are in fact integral parts of the protein complexes at MCSs. We first determined whether TMED2/10 interact with known components of ER-Golgi MCSs by performing co-immunoprecipitation experiments. The immunoprecipitation of endogenous OSBP pulled down both TMED2 and 10 (Figure 5A, B).

**Figure 5.**
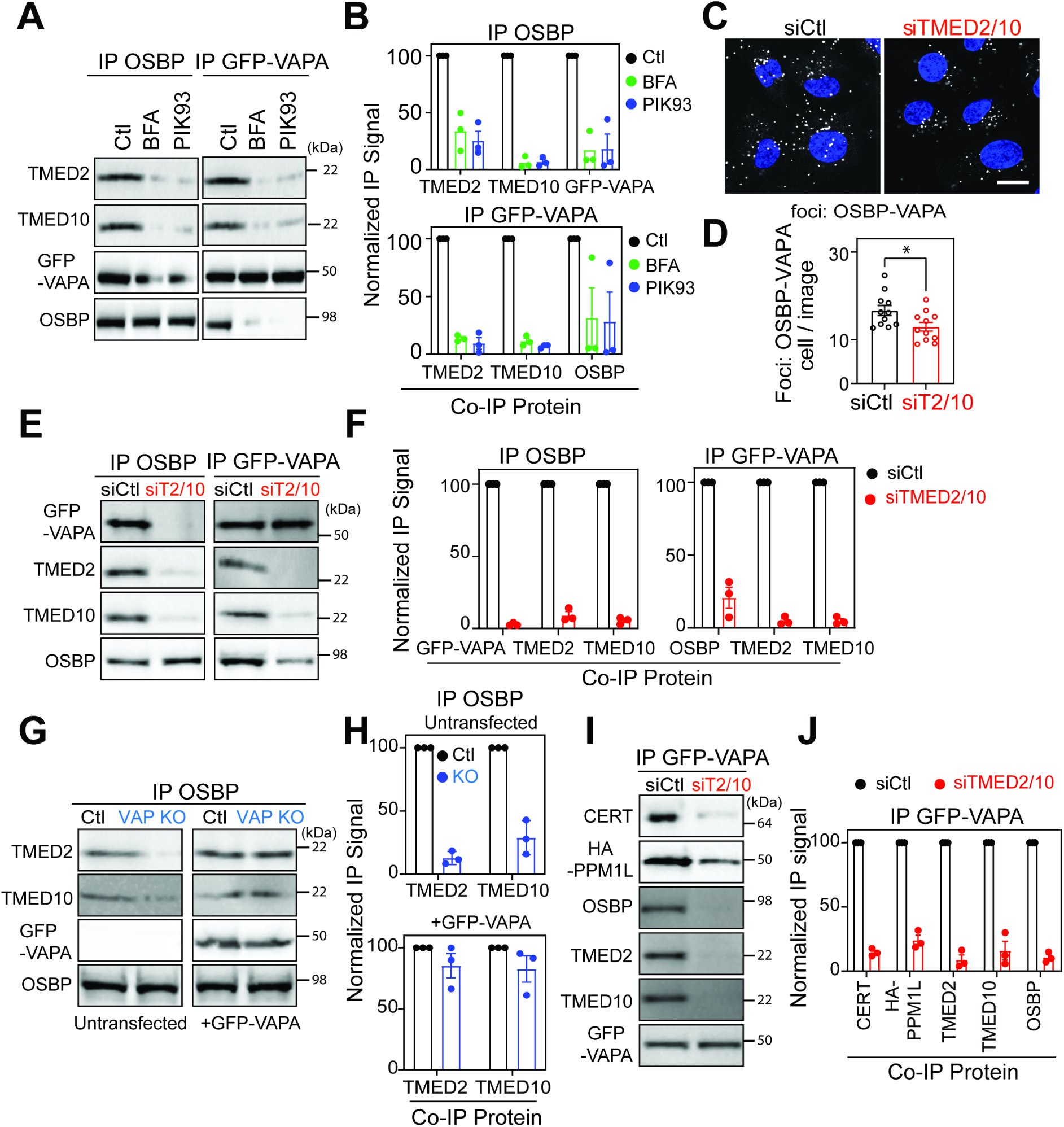
TMED2/10 interact with ER-Golgi MCS proteins and control PtdIns(4)*P* transfer. See also Figure S5. **A)** Western blot of immunoprecipitated (IP) OSBP or IP GFP-VAPA fractions after BFA or PIK93 treatment. Cells were lysed post BFA (5 µg/ml for 5 min) or PIK93 (4 µM for 10 min) treatments, and subsequent IP was performed. **B)** The values for (A) were normalized, with the control set to 100. **C)** Antibody-based *in situ* proximity ligation assay was performed on pre-fixed and permeabilized siCtl and siTMED2/10 cells. Plus and Minus probes were used to target rabbit anti-OSBP and mouse anti-VAPA antibodies. **D)** The number of foci were extracted per image in (C) and are shown here normalised to the number of cells per image. n = 129, siCtl; n = 132, siTMED2/10. Scale bar is 10 µm. *p < 0.05 (Unpaired two-tailed t-test). **E)** and **F)** Same as in (A) and (B) IP OSBP and IP GFP-VAPA was performed in control and siTMED2/10 cells. (F) The Co-IP signal was set to 100 in siCtl cells. **G)** and **H)** Same as in (E and F) IP GFP-VAPA was performed in cells overexpressing HA-PPM1L or GFP-CERT. **I)** and **J)** Same as in (G and H) in HeLa control or VAP knockout cells either transfected with GFP-VAPA or not (untransfected). (J) The Co-IP signal was set to 100 in control cells. Results are mean ± SEM (n ≥ 3).

Since the interaction of OSBP with the Golgi depends on the presence of ARF1 and PtdIns(4)*P*, we tested the effect of the fungal metabolite brefeldin-A (BFA) that inhibits ARF activation, and of the PI4P-kinase inhibitor PIK93 on the TMED2/10-OSBP interaction (Donaldson et al., 2005; Hsu et al., 2010). Our treatment times were chosen based on the release of OSBP from the Golgi (Figure S5A). Immunoprecipitation of OSBP after BFA (5 min) or PIK93 (10 min) treatment showed loss of OSBP-TMED interaction.

Immunoprecipitation of GFP-VAPA also pulled down both TMED2 and 10 and showed that these interactions are impaired upon BFA and PIK93 treatments (Figure 5C, D). These observations indicate that VAPA, OSBP, and TMED2/10 are part of a complex, which depends on the presence of PtdIns(4)*P* and ARF1.

We next tested whether TMED2/10 play a role in the formation or maintenance of ER-Golgi MCS by monitoring the interaction of VAPA with OSBP using a proximity ligation assay and found it to be significantly decreased in TMED2/10-silenced cells (Figure 5C, D), despite the high level of background of this assay. Immunoprecipitation of OSBP from TMED2/10 silencing cells completely failed to pull down GFP-VAPA, and *vice-versa*, immunoprecipitation of GFP-VAPA failed to pull down OSBP (Figure 5E, F). We also tested whether the interaction between OSBP and TMED2/10 involves VAPA. Immunoprecipitation of OSBP from VAPA/B knockout cells failed to pull down TMED2 and 10, and the interaction could be restored by re-complementing the cells with GFP-VAPA (Figure 5G, H).

Given our observations on the delayed production of sphingomyelin, we extended our biochemical analyses to include CERT and its phosphatase PPM1L. Immunoprecipitation of GFP-VAPA pulled down both CERT and phosphatase PPM1L in control cells, as expected, but this interaction was drastically reduced upon TMED2/10 silencing (Figure 5I, J). The interactions between VAPA, OSBP, and CERT could not be rescued by treating cells with the ACAT inhibitor Avasimibe (Figure S5B, C), suggesting that it is the physical interactions between the proteins and not the Golgi integrity that maintain the complex in the presence of TMED2/10.

MCS between the ER and the Golgi are well known to have two classes of complexes, one involved in cholesterol transfer and containing VAP-A, SAC1, OSBP and ARF1, and another involved in ceramide transfer, containing VAP-A, PPM1L, CERT. The above analysis shows not only that TMED2/10 are present in both these complexes and required for their existence, but also that these two complexes actually can form super-complexes, mediating transfer of both cholesterol and ceramide, again in a manner that depends on TMED2/10 (Figure 6).

**Figure 6.**
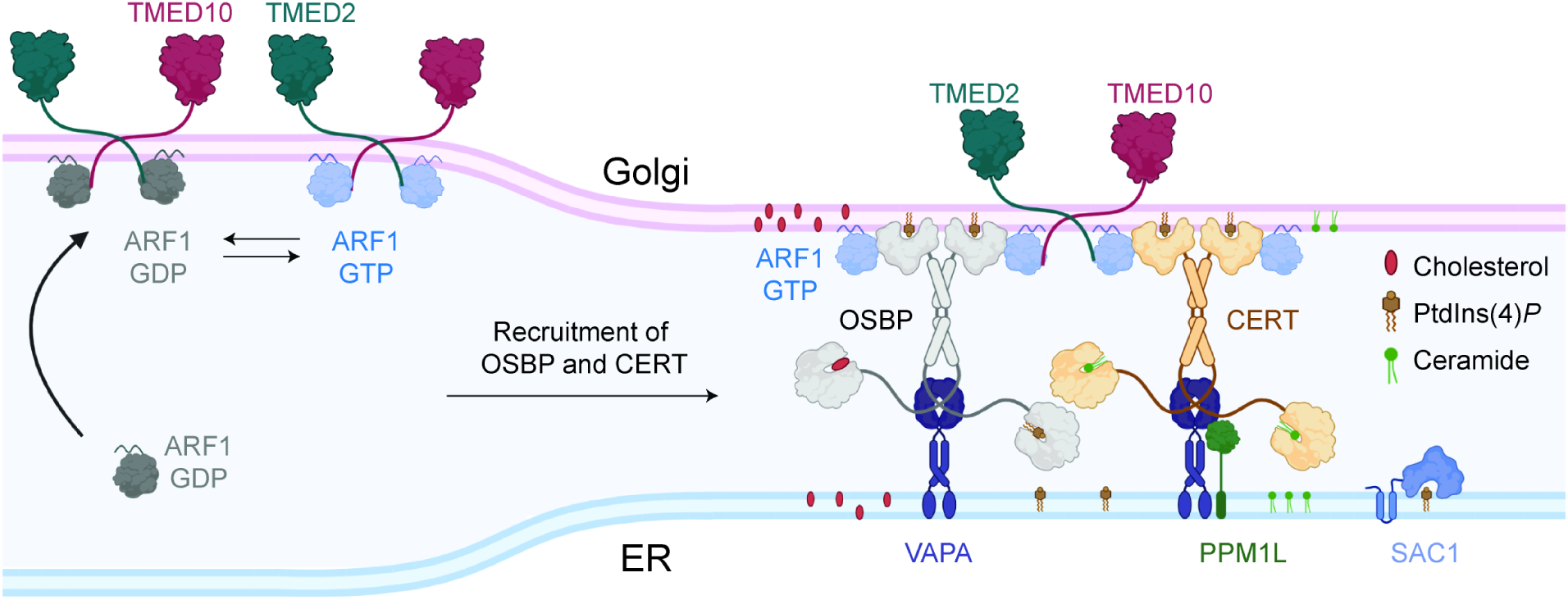
Molecular architecture of ER-Golgi membrane contact sites. Cytosolic ARF1-GDP binds Golgi membranes *via* its interaction with TMED cytosolic tail. Membrane bound ARF1-GDP is then converted to GTP form by the action of a nucleotide exchange factor. ARF1-GTP subsequently recruits OSBP-VAPA and CERT-VAPA complexes to Golgi membranes. Altogether, Golgi localized TMED2/10, ARF1; ER localized VAPA, PPM1L, SAC1 and main transfer proteins OSBP, CERT form a mega-protein structure which constitutes the ER-Golgi MCS. Created with BioRender.com.

## Conclusion

To identify novel factors involved in the anthrax toxin entry process, we herein performed an RNAi screen. We in particular identified two proteins, TMED2 and TMED10 and show that they are involved in plasma membrane sub-compartmentalization. While TMEDs are well established cargo receptors/chaperones of GPI-anchored proteins, we here determined that they are also key players of ER-Golgi MCSs, where they act as indispensable components of large protein super-complexes responsible for intracellular lipid transport that controls the formation of cholesterol-rich plasma membrane nanodomains (Figure 6).

Elegant *in vitro* studies have shown that a VAPA-OSBP complex is sufficient to transfer cholesterol and PtdIns(4)*P in vitro* between vesicles (de la Mora et al., 2021). In living cells, this process in addition requires ARF1 that, by cycling between its GDP and GTP-bound forms, contributes to the recruitment of OSBP at the ER-Golgi MCSs (Adarska et al., 2021; Donaldson et al., 2005; Nakatsu & Kawasaki, 2021). The mechanism that ensures initial recruitment of ARF1 at MCSs is not known. Interestingly, the cytosolic tail of TMED10 has been shown to directly interact with the GDP-bound form of ARF1 and to recruit it to the Golgi (D. U. Gommel, 2001; Zheng et al., 2013). Our work shows that the structural stability, and thereby the function, of the cholesterol transfer complex at ER-Golgi interfaces requires TMED2/10. It is then tempting to speculate that TMED2/10 primes the dynamic assembly the lipid transferring machinery at ER-Golgi MCS by guiding the local recruitment of ARF1 (Figure 6). Future studies will address whether and how the different functions reported of TMED proteins, such as cargo receptors ensuring the transport of the nano-domain associated escorting GPI-anchored proteins (Castillon et al., 2011; Zavodszky & Hegde, 2019) or orchestrating MCSs (the present study), are connected.

Our work also shows that the cholesterol transfer and the ceramide transfer complexes are in fact part of a TMED2/10 dependent supercomplex, which offers interesting potential of co-regulation. The most stunning finding of our work is however that alterations in these supercomplexes can lead to a complete loss of lipid nano-domains at the plasma membrane, and thus that the transport of cholesterol via vesicular trafficking through the biosynthetic pathway is unable to compensate even with time.

ER-Golgi MCSs therefore control plasma membrane compartmentalization, anthrax intoxication and possibly other nanodomain-dependant cellular processes.

## Acknowledgements

We thank Pietro De Camilli Lab (Yale School of Medicine) for VAP KO cells; Hedge Lab (MRC Laboratory of Molecular Biology) for TMED10-HA plasmids; Holthuis Lab (University of Osnabrück) and G. Fairn (Dalhousie University) for D4-mCherry; Y. Hannun and D. Canals (Stony Brooks Cancer Centre) for Lysenin-GFP; A. De Matteis (University of Napoli) for VAPA-GFP plasmid; D. Moreau and S. Vossio (ACCESS Geneva) for high-throughput imaging; J. Paz and T. Hannich for lipidomics; EPFL core facilities BIOP, GECF and BICC for sharing their equipment and expertise; all VDG and D’Angelo Lab members for the discussions; S. Ho for cloning and protein purification; A. Samurkas for protein modelling and N. Panyain for the schematics created with BioRender.com. This work was supported by the Swiss National Science Foundation grant to F.G.v.d.G and to G.D’A.

## Author contributions

M.U.A., O.A.S., L.A., G.D’A. and F.G.v.d.G. conceptualized the overall project, analyzed the results and prepared the manuscript, with input from all co-authors. O.A.S. and I.L. performed the siRNA screen and its analysis, under the guidance of P.L, G.D’A. and F.G.v.d.G. For other experiments, M.U.A., O.A.S., L.A., F. M., T.A., A. C., L.C. performed and analyzed the experiments, with guidance from G.D’A. and F.G.v.d.G.

## Methods

### Cell lines and tissue culture

RPE-1 (CVCL_4388), HeLa (CVCL_0030), and HeLa-MZ (not annotated in CVCL), HeLa-M and HeLa-M VAP KO (Dong et al., 2016) cells were used in this study. RPE1 were grown in DMEM GlutaMAX (Gibco 31966) supplemented with 10% FBS (Pan Biotech) and 2 mM antibiotics (P/S: penicillin and streptomycin). HeLa and HeLa-MZ cells were grown in MEM (Sigma-Aldrich M4655) supplemented with 10% FBS, 2 mM L-glutamine (Gibco 25030081), and 1X MEM Non-Essential Amino Acid Solution (Gibco 11140035) and 2 mM antibiotics (P/S). Experiments were conducted using RPE1 cells, except indicated otherwise.

### Antibodies

Primary antibodies used in this study that are commercially available include: rabbit anti-MEK2 (Santa Cruz Biotechnology sc-523, AB_2281672), mouse anti-EEA1 (BD Biosciences 610457, AB_397830), goat anti-protective antigen from *Bacillus anthracis* (List Biological Laboratories #771B), mouse anti-GAPDH (Sigma-Aldrich G8795, AB_1078991), rabbit anti-CMG2 (Proteintech Group 16723–1-AP, AB_2056741), rabbit anti-OSBP (Atlas antibodies HPA039227, AB_2676401), rabbit anti-CERT (Novus Biologicals NBP2-59018), rabbit anti-CLIMP63 (Bethyl Laboratories A302-257A, AB_1731083), rabbit anti-caveolin1 (Santa Cruz Biotechnology sc-894, AB_2072042), rat anti-HA-HRP (Roche Diagnostics 12013819001, AB_390917), mouse anti-V5 (Thermo Fisher Scientific R960-25, AB_2556564), mouse anti-VAPA (Santa Cruz Biotechnology sc-293278; AB_2801294), mouse anti-TMED2 (Santa Cruz Biotechnology sc-376459; AB_11150297), rabbit anti-TMED10 (Bethyl Laboratories A305-219A; AB_2631612), rabbit anti-GOLPH3 (Abcam ab98023; AB_10860828), mouse anti-P230 (BD Biosciences 611281; AB_398809), mouse anti-GM130 (BD Biosciences 610823; AB_398142), and rabbit anti-Giantin (Biolegend 94302), mouse anti-GFP (Roche AB_390913), rabbit anti-CERT (Abcam ab151285).

Primary antibodies that were home-made include: anti-MEK1 raised in rabbit (Abrami et al., 2003) and anit-flotillin1 raised in rabbit (Abrami et al., 2008). Secondary antibodies for Western blotting include anti-mouse-HRP (GE Healthcare NA931, AB_772210), anti-rabbit-HRP (GE Healthcare NA934, AB_772206), anti-goat-HRP (Sigma-Aldrich A5420, AB_258242). For immunofluorescence, secondary antibodies used include goat anti-mouse Alexa Fluor 488 (Thermo Fisher Scientific A11029; AB_2534088) and donkey anti-rabbit Alexa Fluor 568 (Thermo Fisher Scientific A10042; AB_2534017).

### Toxins

Anthrax toxin was purified out of *E. coli* as described previously (Sergeeva & van der Goot, 2019). Other toxins used in this study were kind gifts from different labs. These include, D4-mCherry (Maekawa & Fairn, 2015) and Lysenin-GFP (Canals et al., 2018).

### Gene Silencing and Overexpression

Genes were silenced using siRNAs obtained from Qiagen (see Table S2 for sequences). Silencing was performed for 72–96 h using Lipofectamine RNAiMAX (Thermo Fisher Scientific 13778150) or INTERFERrin (Polyplus 409-10) following the manufacturer’s protocol. Silencing efficiency was checked via Western blot and/or qPCR. TMED10-HA constructs were a kind gift from Prof. Ramanujan Hedge, MRC laboratory, UK. Proteins were expressed in RPE1 or HeLa cells for 24–48 h using TransIT®-LT1 Transfection Reagent (Mirus Bio) or Neon Electroporation (ThermoFisher) following the manufacturer’s protocol or ROTI Fect (Carl Roth).

### Screening

The siRNA screen consisted of the four 384-well plates used in the Endocytome screens (Liberali et al., 2014) and one extra 384-well plate with genes that code for surface proteins. The extra plate was designed using cell surface mass spectrometry data in RPE1, the cell surface protein atlas (Bausch-Fluck et al., 2015) and known surface GPI-anchored proteins. Plates were ordered with pools of three Ambion Silencer Select siRNAs per gene (Thermo Fisher Scientific). The screen used reverse transfection of siRNAs in which all wells had 250 nM siRNA concentration in 10 µL of Opti-MEM (Gibco; Thermo Fisher Scientific). Lipofectamine2000 (Thermo Fisher Scientific) transfection reagent was added in at 0.05 µL in 5 µL of Opti-MEM per well.

Cells were plated into a transfection mix at 800 per well in 50 µL of complete media. After 72 hours of growth, cells were treated with anthrax toxin (0.5 µg/mL PA and 0.1 µg/mL LF) in minimal media (Glasgow minimal essential media (Sigma-Aldrich G6148) buffered with 10 mM HEPES) and incubated at 4 °C for 1 h. Wells were washed twice with complete media and plates were shifted to 37 °C for 1.5 h. After which, wells were fixed with 4% PFA in PBS for 15 min, quenched with 50 mM NH_4_Cl in PBS for 10 min, permeabilized with 0.1% TritonX in PBS for 5 min, with 2 PBS washes between all steps. Next, cells were blocked with 0.5% BSA in PBS for 30 min and primary antibodies (rabbit anti-MEK2N 1:100 and mouse anti-EEA1 1:200) were applied in blocking buffer for 1 h, after which secondary antibodies (Alexa488 anti-rabbit and Alexa568 anti-mouse both at 1:600) were applied in blocking buffer for 45 min. Finally, DAPI was applied at 0.4 µg/mL final in PBS for 10 min and CellTrace (Alexa 647 at 1:5000) in carbonate buffer was applied for 5 min. Wells were washed with PBS twice between steps and kept in PBS for imaging. Imaging was done using the Yokogawa CV7000S using the 20X objective and imaging the full well. Z-stack sums for each image were stitched together for the full well image.

### Screen segmentation and quantification

The wells/siRNAs of the screens were quantified using CellProfiler (Kamentsky et al., 2011). Briefly, the images were separated into the four channels: DAPI, MEK2N, CellTrace, and EEA1. For the toxin entry, looking at MEK2 cleavage and therefore, disappearance of MEK2N, the quantification was done as follows. The cells were selected using the nuclei in the DAPI channel. Then, the cells were segmented using the CellTrace channel. To only quantify the cytoplasm, a mask for the cell was done excluding the nucleus. Finally, the median intensity of both MEK2N and CellTrace of each cell was calculated, and from that, the mean intensity of both MEK2N and CellTrace of all the cells in each well was computed. For each well, and therefore siRNA, both mean intensities were converted to 16bit. The MEK2N (0-1) and CellTrace (0.01-1) for each well were normalized per plate and the MEK2N value was divided by CellTrace to have a ratio of remaining MEK2. Finally, the Z-score was calculated of this ratio for each siRNA per plate. As there were duplicates in the screen, some later analyses used the average of the two Z-scores as the value for that well and siRNA. For the EEA1 screen, the images in the EEA1 channel were thresholded to remove the highest and lowest 0.05% pixels. The total perimeter of that channel was quantified, and then normalized by dividing by the square of the log base 10 of the cell number of each well. This value was then Z-score normalized per plate. As with the Toxin Entry screen, averages of the two Z-scores for the EEA1 screen were computed for later analyses. For both screens, a few siRNAs were removed that either were missing images or had less than 100 cells in their wells.

### SDS/PAGE and Western Blots

Cells were normally lysed in immunoprecipitation buffer [IPB: 0.5% Nonidet P-40, 500 mM Tris pH 7.4, 20 mM EDTA, 10 mM NaF, 2 mM benzamidin, and a Roche mini-protease inhibitor mixture tablet (PI tab; Sigma-Aldrich 11836153001) and spun at 5,000 × g for 3 min to eliminate the DNA. Exceptionally, cells transfected with the PC- sensors were lysed in PBS supplemented with 1% Triton X-100, 1 mM EDTA, and 1 PI tab, and spun at 21,000 × g for 10 min. DNA-free lysates were quantified using Bicinchoninic acid (BCA) protein assay (Interchim Uptima 40840A) and denatured by addition of SDS sample buffer with β-mercaptoethanol and incubation for 5–10 min at 95 °C. Samples were migrated on precast Novex 4–20% or 4–12% polyacrylamide gels (Thermo Fisher Scientific), then transferred to Novex nitrocellulose membranes (Thermo Fisher Scientific) using iBlot 2. Blocking and antibody steps were performed using 5% milk in PBST (PBS with 0.5% Tween-20). Primary antibody steps were incubated overnight at 4 °C, while the membranes were incubated with secondary antibodies for 1 h at room temperature, both with gentle shaking. Three to five washes of PBST were performed before developing using the Fusion Solo-S (Viber-Smart Imaging) and the Fusion Solo chemiluminescence imaging system.

### Anthrax toxin entry assays

Cells at 80–90% confluence were washed two times with minimal media [Glasgow minimal essential media (Sigma-Aldrich G6148) buffered with 10 mM HEPES] at 4 °C and then incubated with toxin (anthrax toxin: 500 ng/mL PA and 50 ng/mL LF) for 30 min to 1 h at 4 °C. For cleaved (or pre-nicked) toxin, PA was incubated for 10 min with 100 μg/mL trypsin then stopped with 1 mg/mL of trypsin inhibitor for 1 min, and added to minimal media as for the uncleaved toxin. After the toxin incubation, cells were washed three times with minimal media and moved to 37 °C. The cells were lysed at the appropriate times shown in the experiments (0–2.5 h for experiments with anthrax toxin showing MEK cleavage; 0–90 min for experiments showing PA cleavage). For acid pulse treatment, lysates were treated with a 10% addition of isotonic buffer (145 mM NaCl, 20 mM MES-Tris pH 4.5) for 10 min at RT. Samples were run on gels as described above with GAPDH as loading controls. For comparison purposes, quantified values were normalized to time 0 (for MEK experiments) or within each experiment (for PA experiments). The molecular weight of the SDS-resistant PA^oligo^ is not defined as it is above the top marker in SDS/PAGE. Experiments were repeated at least three times and representative blots are shown in the figures.

### Cell Surface Protein Pull-Down

Surface biotinylation assays were performed as described previously (Sergeeva & van der Goot, 2019) . Briefly, cells were shifted to ice and incubated for 30 min with cold biotinylation solution (Thermo Fisher Scientific EZ-link Sulfo-NHS 21327) diluted in PBS (0.17-mg/mL final concentration), then quenched three times with cold 100 mM NH4Cl. Cells were lysed as normally and about 5% of the DNA-cleared lysate was taken as a TCE control. The rest of the lysate was incubated with prewashed streptavidin-coupled beads (Sigma S1638) overnight at 4 °C. Finally, beads were washed and proteins were eluted from beads as above. GAPDH was used as an intracellular control. Experiments were repeated at least three times and representative blots are shown in the figures.

### Lipid extraction

Total lipid extracts were prepared using a standard MTBE protocol for total lipid analysis by mass spectrometry. Briefly, cell pellet was resuspended in 100 μL H_2_O. 360 μL methanol and 1.2 mL of MTBE were added and samples were placed for 10 min on a vortex at 4 °C followed by incubation for 1 h at room temperature on a shaker. Phase separation was induced by addition of 200 μL of H2O. After 10 min at room temperature, samples were centrifuged at 1000x g for 10 min. The upper (organic) phase was transferred into a glass tube and the lower phase was re-extracted with 400 μL artificial upper phase [MTBE/methanol/H_2_O (10:3:1.5, v/v/v)]. The combined organic phases were dried in a vacuum concentrator. The dried lipid pellet was then extracted using 300 μL water-saturated n-butanol and 150 μL H_2_O. The organic phase was collected, and the aqueous phase was re-extracted twice with 300 μL water-saturated n-butanol. The organic phases were pooled and dried in a vacuum concentrator.

### LC-MS untargeted lipidomics

For phospholipid analysis, lipid extracts (2 μL injection volume in CHCl_3_:MeOH 2:1) were separated over an 8 min gradient at a flow rate of 200 μL/min on a HILIC Kinetex Column (2.6lm, 2.1 × 50 mm^2^) on a Shimadzu Prominence UFPLC xr system (Tokyo, Japan). Mobile phase A was acetonitrile:methanol 10:1 (v/v) containing 10 mM ammonium formate and 0.5% formic acid while mobile phase B was deionized water containing 10 mM ammonium formate and 0.5% formic acid. The elution of the gradient began with 5% B at a 200 μL/min flow and increased linearly to 50% B over 7 min, then the elution continued at 50% B for 1.5 min and finally, the column was re-equilibrated for 2.5 min. MS data were acquired in full-scan mode at high resolution on a hybrid Orbitrap Elite (Thermo Fisher Scientific, Bremen, Germany). The system was operated at 240,000 resolution (*m/z* 400) with an AGC set at 1.0E6 and one microscan set at 10-ms maximum injection time. The heated electrospray source HESI II was operated in positive mode at a temperature of 90 °C and a source voltage at 4.0KV. Sheath gas and auxiliary gas were set at 20 and 5 arbitrary units, respectively, while the transfer capillary temperature was set to 275 °C.

Mass spectrometry data were acquired with LTQ Tuneplus2.7SP2 and treated with Xcalibur 4.0QF2 (Thermo Fisher Scientific). Lipid identification was carried out with Lipid Data Analyzer II (LDA v. 2.6.3, IGB-TUG Graz University). The LDA algorithm identifies peaks by their respective retention time, m/z and intensity. Care was taken to calibrate the instrument regularly to ensure a mass accuracy consistently lower than 3 ppm thereby leaving only few theoretical possibilities for elemental assignment.

### Sphingosine Pulse and Chase

RPE1 cells were pulse-labelled in serum-free 2% fatty-acid-free BSA in Dulbecco’s modified Eagle’s medium, at a final concentration of 1 μCi/mL [3H]-sphingosine, for 2 h. Cells were chased in complete medium for the indicated times, harvested and processed for lipid extraction. Lipids were spotted onto silica-gel high performance-TLC (HPTLC) plates (Merck, Germany)(Capasso et al., 2017), and resolved with a mixture of chloroform: methanol: water (65:25:4 v/v/v). To visualise the unlabelled standards (i.e., Cer, GlcCer, LacCer, Gb3, SM and GM3) the TLC plates were placed in a sealed tank saturated with iodine vapour, while the radiolabelled lipids were analysed using a RITA**^®^** TLC Analyser (Raytest, Germany), and quantified using GINA**^®^** (Raytest, Germany) software analysis.

### Cholesterol Measurement

Cholesterol and cholesterol esters were measured using Amplex Red Cholesterol Assay kit (Thermofisher, MAK043) as per the manufacturer’s protocol.

### Detergent Resistant Membrane (DRMs)

DRMs were prepared from RPE1 cells as described previously using OptiPrep (Alere Technologies #1114542) gradients as previously published (Abrami et al., 2003; Sergeeva & van der Goot, 2019). Briefly, cells were lysed on ice in TNE buffer (25 mM TrisHCl, pH 7.4; 150 mM NaCl; 5 mM EDTA) supplemented with 1% Triton X-100 and applied to the bottom of a gradient consisting of three layers: 100% OptiPrep, 50% OptiPrep and 50% TNE, and 100% TNE. After a 2-h run at 55,000 rpm in a Thermo Fisher Scientific S55-S rotor, six equal fractions were carefully collected from the top. Fractions were run on SDS/PAGE for western blots or used for mass spectrometry. Experiments were repeated at least three times and representative blots are shown in the figures.

### Mass spectrometry

SDS-PAGE gel slices were washed twice in 50% ethanol and 50 mM ammonium bicarbonate for 20 min and dried by vacuum centrifugation. Samples reduction was performed with 10 mM dithioerythritol for 1 h at 56 °C. A washing-drying step as described above was repeated before performing sample alkylation with 55 mM Iodoacetamide for 45 min at 37 °C in the dark. Samples were washed-dried again and digested overnight at 37 °C using Mass Spectrometry grade trypsin at a concentration of 12.5 ng/µL in 50 mM ammonium bicarbonate and 10 mM calcium chloride. Resulting peptides were extracted in 70% ethanol, 5% formic acid twice for 20 min with permanent shaking. Samples were further dried by vacuum centrifugation and stored at -20 °C. Peptides were desalted on SDB-RPS StageTips (Rappsilber et al., 2007) and dried by vacuum centrifugation. For TMT labelling, peptides were first reconstituted in 8 μL HEPES 100 mM (pH 8.5) containing 10 ng trypsin-digested Chicken Ovalbumin. Labeling was performed by adding 3 μL of TMT solution (20 µg/μL in pure acetonitrile) and incubating samples at room temperature for 1.5 h. Reactions were quenched with hydroxylamine to a final concentration of 0.4% (v/v) for 15 min. TMT-labelled samples were then pooled at a 1:1 ratio across all samples and dried by vacuum centrifugation. Samples were then fractionated into 12 fractions using an Agilent OFF-GEL 3100 system. Resulting fractions were desalted on SDB-RPS StageTips and dried by vacuum centrifugation. Each individual fraction was resuspended in 10 μL of 2% acetonitrile, 0.1% formic acid and nano-flow separations were performed on a Dionex Ultimate 3000 RSLC nano UPLC system on-line connected with a Lumos Fusion Orbitrap Mass Spectrometer. A capillary precolumn (Acclaim Pepmap C18, 3 μm-100 Å, 2 cm x 75 μm ID) was used for sample trapping and cleaning. Analytical separations were performed at 250 nL/min over 150 min. biphasic gradient on a 50cm long in-house packed capillary column (75μm ID; ReproSil-Pur C18-AQ 1.9μm; Dr. Maisch). Acquisitions were performed through Top Speed Data-Dependent acquisition mode using a 3 seconds cycle time. First MS scans were acquired at a resolution of 120,000 (at 200 m/z) and the most intense parent ions were selected and fragmented by High energy Collision Dissociation (HCD) with a Normalized Collision Energy (NCE) of 37.5% using an isolation window of 0.7 m/z. Fragmented ions were acquired with a resolution 50,000 (at 200 m/z) and selected ions were then excluded for the following 120 s.

Raw data were processed using SEQUEST, Mascot, MS Amanda (Dorfer et al., 2014) and MS Fragger (Kong et al., 2017) in Proteome Discoverer v.2.4 against a concatenated database consisting of the Uniprot Human Reference Proteome (Uniprot Release: 2019_06) and common contaminants including the Ovalbumin from chicken (Uniprot Accession Number: P01012). Enzyme specificity was set to Trypsin and a minimum of six amino acids was required for peptide identification. Up to two missed cleavages were allowed. A 1% FDR cut-off was applied both at peptide and protein identification levels. For the database search, carbamidomethylation (C), TMT tags (K and Peptide N termini) were set as fixed modifications, while oxidation (M) was considered as a variable one. Resulting text files were processed through in-house written R scripts (version 3.6.3) (Schindelin et al., 2012). The unnormalized abundances calculated by Proteome Discoverer were transformed in log2 and subtracted to obtain ratios. The Z-scores of the ratio of ratios [F2 (siTMED2/10 / siCtl) / Input (siTMED2/10 / siCtl)] were calculated.

### Immunofluorescence

HeLa or RPE1 cells were plated at about 50% confluency after transfection. Cells were washed with PBS, fixed in 4% paraformaldehyde (PFA), washed thoroughly in PBS, quenched with 50 mM NH_4_Cl, and washed again with PBS. When necessary, cells were permeabilized with 0.1% Triton X-100 in PBS for 5 min. After, cells were washed and blocked with 1% BSA in PBS. Primary and secondary antibody incubations were done in the same buffer. Finally, coverslips were mounted in Prolong Glass Antifade Mountant (ThermoFisher Scientific P36980). Images were acquired using either confocal LSM710 (Carl-Zeiss AG) or Leica-SP8 (Leica Biosystems) with 63X objective and a pinhole size of 1 AU and at least 2 line-averaging. Images were processed using Fiji ImageJ (Schindelin et al., 2012).

### D4 staining

Cells were prefixed with PFA, blocked and stained with home purified D4-mCherry for 1 h at room temperature. Coverslips were mounted on glass slides and images were acquired as mentioned previously.

### Lysenin staining

Cells were starved in FBS free medium overnight before incubation with Lysenin-GFP for 30 min at 4° C. Cells were then fixed using PFA for 10 min, mounted and imaged as explained above.

### Automated fluorescence microscopy and quantification

Cells were plated at 2x10^4^ cells per well and silenced for 72h with either siCtl and siTMED2/10 following manufacturer’s instructions in Ibidi 96-well µplates. Cells were fixed with 3% PFA and permeabilized with 0.05% saponin. Cells were then stained with primary antibodies followed by incubation with (when required) secondary antibodies tagged with alexa-fluor (488 or 568) with Hoechst. Images were acquired with either 40X or 60X plan Apo.

MetaXpress Custom Module editor software from Molecular Devices was used to extract different parameters such as signal intensity or objects, as in previous publications (Larios et al., 2020; Moreau et al., 2019). Cell mask was created using Hoechst to create the master object (cell). Golgi was segmented using either GM130 or GOLPH3. These masks were then applied on the fluorescent images to extract relevant measurements. When indicated, a perinuclear mask (4-μm-diameter region immediately outside the nucleus) to quantify cholesterol (filipin) signals inside and outside this perinuclear area was used. The final masks were applied to all original fluorescent images and measurements per cell and averages per well were extracted. For GM130-positive objects, all objects per cell were analysed. The same analysis pipeline was applied to all images.

### In situ Proximity Ligation Assay

*In situ* proximity ligation (PLA) assay were performed using materials and guidelines provided with DuoLink PLA kit (Sigma-Aldrich, USA). Briefly, cells were fixed and permeabilized as mentioned above. Cells were blocked in DuoLink blocking solution and incubated with rabbit anti-OSBP and mouse anti-VAPA antibodies. Cells were next incubated with anti-mouse PLUS and anti-rabbit MINUS secondary probes followed by ligation and amplification with specific reagents. Cells were imaged using a laser-scanning-confocal microscope. Images were processed using Fiji ImageJ and PLA spots were quantified using TrackMate (Schindelin et al., 2012).

### Quantitative Real-Time PCR

RNA was extracted from RPE1 cells using the Qiagen RNAeasy kit with use of the QIAshredder. RNA concentration was measured and 500 ng of total RNA was used for cDNA synthesis using iScript (Bio-Rad 1708891). A 1:5 dilution of cDNA was used to perform quantitative real-time PCR using Applied Biosystems SYBR Green Master Mix (Thermo Fisher Scientific) on 7900 HT Fast QPCR System (Applied Biosystems) or QuantStudio 6 Pro Real-Time PCR System (Thermo Fisher Scientific) with SDS 2.4 Software. Primers (see Table S2 for sequences) were validated with standard curves over 4 dilutions. The data (always in triplicate) were normalized using multiple housekeeping genes (TATA-binding protein, β-microglobulin and β-glucuronidase, 5’-aminolevulinic acid synthase 1). For real-time quantitative PCR, data was processed using R (R Core Team, 2019).

### RNA sequencing

RNA quality was controlled on the TapeStation 4200 (Agilent), confirming that all were of good quality (scores >9.6). Libraries for mRNA-seq were prepared with the Stranded mRNA Ligation method (Illumina) starting from 1ug RNA, according to manufacturer’s instructions. Libraries, all bearing unique dual indexes, were subsequently loaded at 1.44 pM on a NextSeq 500 high output flow cell (Illumina) and sequenced according to manufacturer instructions, yielding pairs of 80 nucleotides reads. Reads were trimmed of their adapters with bcl2fastq v2.20 (Illumina) and quality-controlled with fastQC v0.11.9.

Paired-end sequencing reads were aligned using STAR (version 2.7.9a) to the Human reference genome GRCh38 (gencode v36, Ensembl 102). Quantification of uniquely mapped reads to the genes was performed with HTSeq counts (Part of the ’HTSeq’ framework, version 0.12.4) with the following parameters *-s reverse -m intersection-nonempty -r pos -t exon -i gene_id -q*. Counts pre-processing and differential analysis was done in R (version 4.0), using edgeR (v. 3.30.3) and DESeq2 package (v. 1.28.1)(Robinson et al., 2010). A total of 11’995 protein-coding genes with a cpm value greater than 1 in at least 2 samples were considered for the rest of the analysis.

### Coimmunoprecipitations

Coimmunoprecipitations were performed as described previously (Bürgi et al., 2020; Sergeeva & van der Goot, 2019). Briefly, RPE1 or HeLa cells were lysed normally as described above. A tenth of the lysate was taken as TCE control and the rest was added to washed Protein G-coupled Sepharose beads (Sigma-Aldrich GE Healthcare 17-0618-01) for preclearing (30 min). The lysates were then added to freshly washed Protein G beads containing antibody for an overnight incubation at 4 °C. Proteins were eluted off the beads and divided for Western blots for direct immunoprecipitation and coimmunoprecipitation. Experiments were repeated at least three times and representative blots are shown in the figures.

### Quantification, Statistics, Figures

Western blot and lipid droplet quantifications were done using Fiji (Schindelin et al., 2012). Figures were generated and statistics were performed as described in the figure legends using GraphPad Prism or R statistical computing environment (R Core Team, 2019). Throughout the study, indicated significance asterisks are as follows: *P < 0.05, **P < 0.01, ***P < 0.001 and ****P < 0.0001, while unmarked comparisons with control are not significant. All error bars are SEMs. For all areas under curve (AUC) t-testing compared with the control, specific P values are reported on each graph. The pathway analysis figure was made using the ClueGo application of Cytoscape v3.7 (Bindea et al., 2013).

**Table S1: Screen Z-scores.**

Average Z-scores of the two replicates of the Toxin Entry and EEA1 screen are displayed for each silenced gene.

**Table S2: Hit List Sequences (real-time qPCR primer sequences and their amplification efficiencies, siRNA sequences and their knockdown efficiencies).**

The list of genes identified as hits that were subjected to further validation are listed with their gene expression levels in RPE-1 cells (RNA-Seq; TPM) (Sergeeva & van der Goot, 2019), their qPCR primer sequences, the amplification efficiencies of these real-time qPCR sequences (over 4 log2 units), their siRNAs and their knockdown amount upon silencing as read out by qPCR.

---

**Figure S1:**
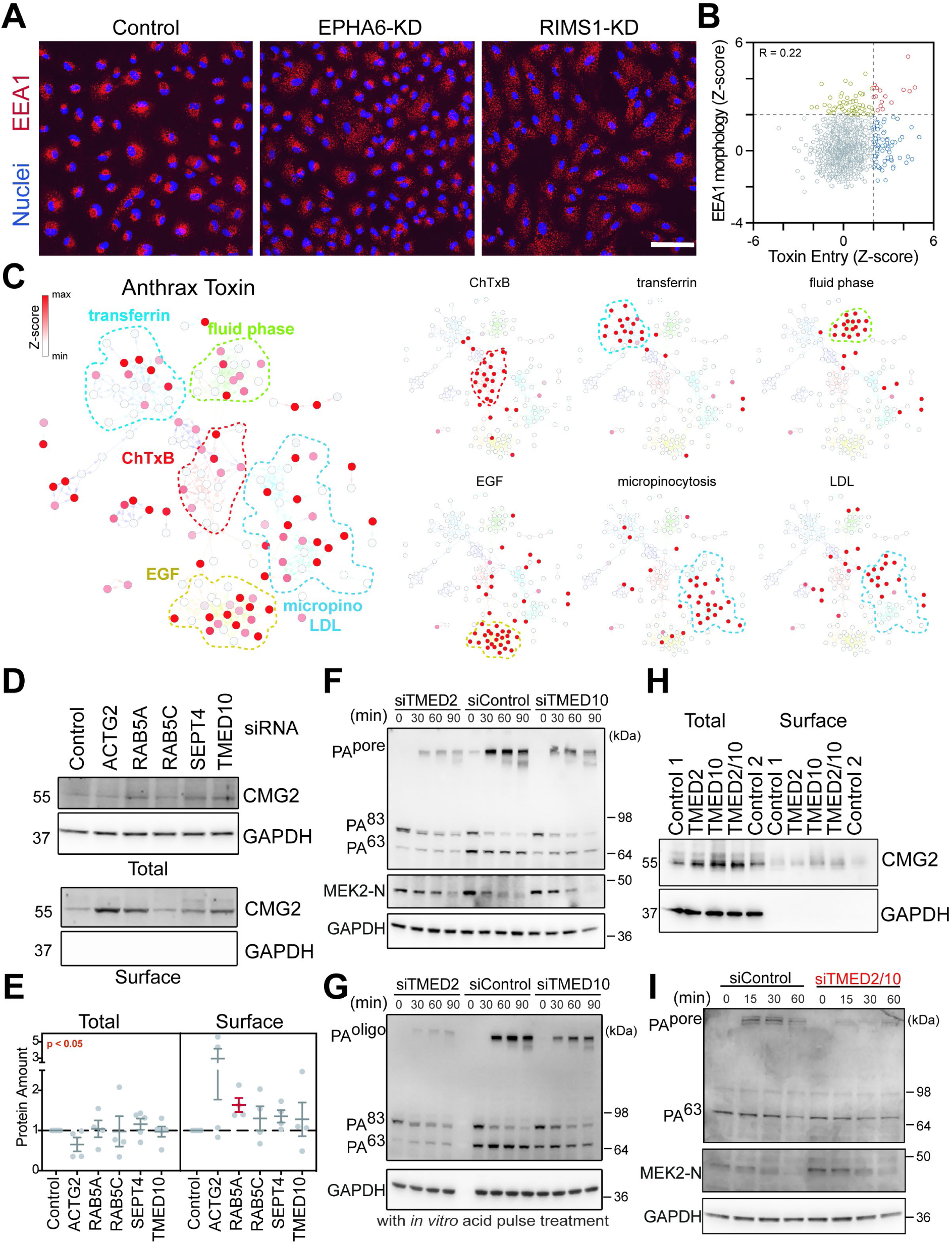
Anthrax toxin entry screen. **A)** Representative IF images obtained in the image-based screen for endosome morphology. DAPI (blue); anti EEA1 antibody (red). Scale bar is 100 µm. **B)** Scatter plot of Z-scores associated with toxin entry *vs.* endosome morphology for each screened gene. Blue represents conditions where anthrax toxin entry is significantly and specifically impaired, yellow represents conditions where endosome morphology is significantly and specifically impaired, and red represents conditions where both anthrax toxin entry and endosome morphology are significantly impaired. **C)** Network visualisation of genes (132) involved in endocytic modules as in Liberali et al, (2014). Genes are colored according to the Z-score associated with the effect of their silencing in the anthrax toxin screen (left panel) and in the screens for different endocytic pathways (right panels). ChTxB, Cholera Toxin uptake; Transferrin; Transferrin uptake; fluid phase, 10kDa Dextran uptake in unstimulated cells; EGF, EGF (50 ng/mL) uptake Macropynocytosis, 10 kDa Dextran uptake in EGF stimulated cells; LDL uptake. Endocytic modules are highlighted by dotted lines coloured according to the most affected pathway. **D)** Cells were biotinylated for 30 min at 4°C, quenched, and lysed. Biotinylated proteins were pulled down with streptavidin beads. Both, total cell extract (TCE) and pull-down fractions were run on SDS-PAGE. Membranes were probed for the anthrax toxin receptor, CMG2, and GAPDH. **E)** Quantification of the total (left) and cell surface (right) levels of CMG2 in the western blot in (D). **F)** Control or silenced cells were treated with (0.5 µg/mL PA83 and 0.1 µg/mL LF) for 1 h at 4 °C, then shifted to 37 °C for multiple time points before lysis. Samples were run on SDS-PAGE. Membranes which were probed with antibodies against PA, MEK2-N and GAPDH. **G)** same as in (F) lysates were treated with an acidic buffer for *in vitro* formation of SDS-resistant pores out of the surface PA oligomers. **H)** Same as in (D). **I)** Same as in (F) cells were silenced for both TMED2/10 and were treated with pre-cleaved PA63. Results are mean ± SEM (n = 4). *p < 0.05 (paired two-tailed t-test).

**Figure S2.**
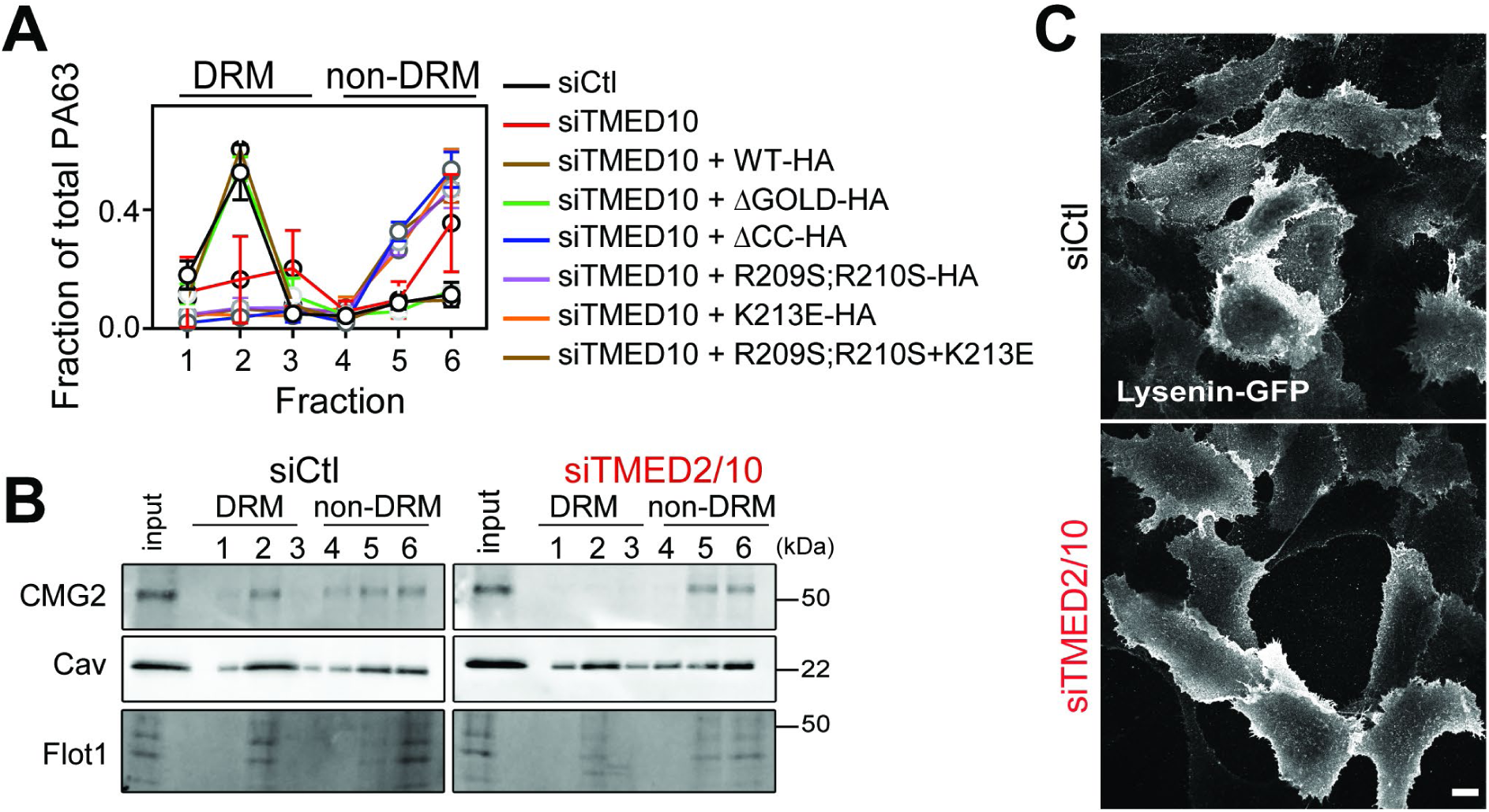
TMED2/10 affects the nanodomain association of CMG2 and Flot1. **A)** (see also Figure 2D). Values were normalised within each experiment. Detergent resistant membranes (DRM) or non- DRM fractions are indicated above each lane. **B)** Step gradient fractions from siCtl and siTMED2/10 cells were run on gel. Membranes were probed with antibodies against CMG2, Caveolin (Cav), and Flotillin1 (Flot1). DRM or non-DRM fractions are indicated above each lane. **C)** Control or silenced cells were stained with Lysenin-GFP and crosslinked with PFA (see methods). Scale bar is 10 µm.

**Figure S3.**
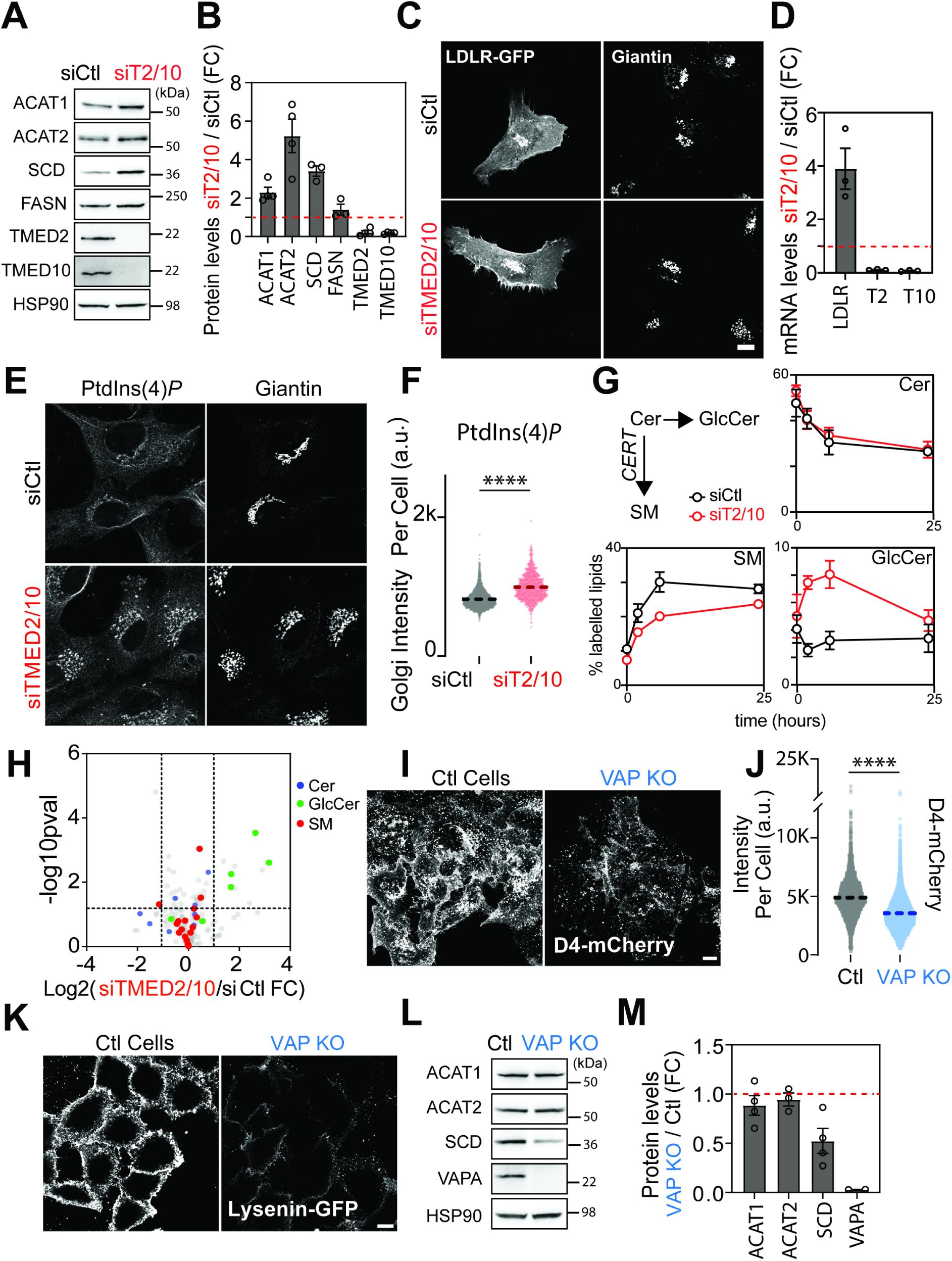
TMED2/10 affects lipid transfer at ER-Golgi MCS. **A)** Western blot of total lysates from siCtl and siTMED2/10 cells. **B)** Protein levels from (A) are expressed as the fold change between siCtl *vs.* siTMED2/10. **C)** LDLR-GFP transfected cells were fixed, permeabilised and stained with Giantin. **D)** RNA was extracted from siCtl and silenced cells 72 h post-transfection and qPCR was performed using specific primers. **E)** siCtl and silenced cells for PtdIns(4)*P* and Giantin. **F)** Golgi mask was created using the Golgi marker. Average PtdIns(4)*P* signal was calculated within that Golgi mask per cell. n = 2467, siCtl; n = 1095, siTMED2/10 from three replicates. **G)** Silenced and control cells were pulse-labelled with ^3^H-labeled shingosine and chased in media for the time points shown. Cells were lysed and lipids were extracted and spotted on HPTLC plates. Plates were scanned using RITA analyzer and values were extracted for each lipid species. **H)** Lipidomic analysis of siTMED2/10 *vs*. siCtl cells. Most important lipid species are highlighted in terms of log2 of the siTMED2/10 over siCtl ratio. **I)** HeLa control and VAP knockout cells were fixed and stained with D4-mCherry and were imaged using confocal microscopy. **J)** The average mCherry signal from (I) was plotted for each cell for both conditions. n = 4327, WT; n = 8369, VAP KO from three replicates. **K)** Control or VAP KO cells were stained with Lysenin-GFP and crosslinked with PFA (see methods). **L and M)** same as (A and B) for HeLa Ctl *vs.* VAP KO cells. Results are mean ± SEM (n ≥ 2). Values were normalized against three house-keeping genes. ****p < 0.0001 (Unpaired two-tailed t-test). All scale bars are 10 µm.

**Figure S4.**
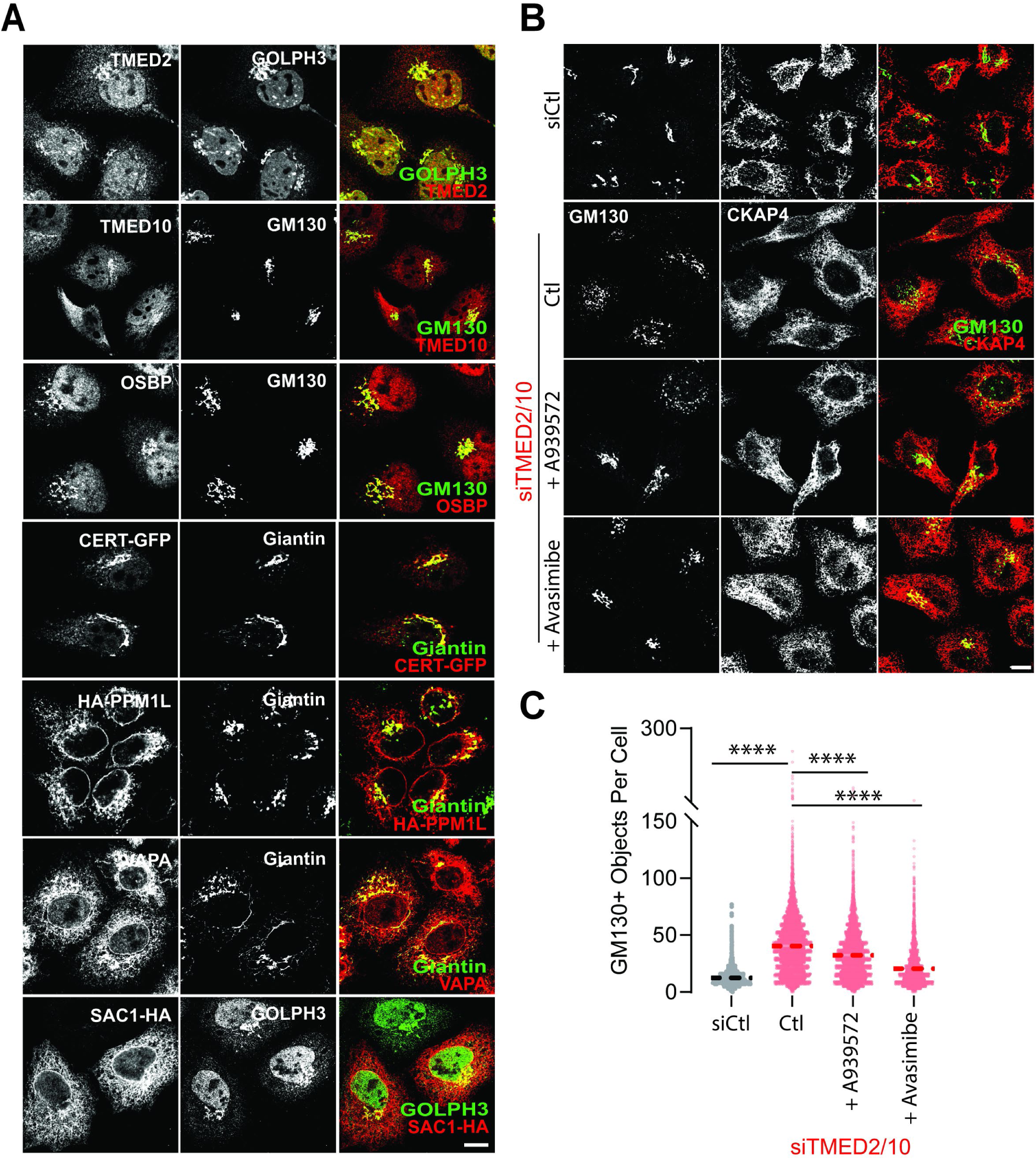
SCD and ACAT inhibition recover Golgi morphology in siTMED2/10 cells. **A)** HeLa cells (siCtl and siTMED2/10) were fixed, permeabilized, and stained for different MCS proteins and Golgi markers. Tagged versions of PPM1L, CERT, and SAC1 were overexpressed 24 h pre-fixation. Images were acquired using laser confocal microscopy. **B)** HeLa control or silenced cells were treated with either A939572 or Avasimibe (10 µM each) or not (Ctl) 24 h before fixation and were stained with GM130 (Golgi) and CKAP4 (ER) markers. **C)** Cells in (B) were segmented using nuclear labelling and GM130+ objects were quantified. n = 7466, siCtl; n = 6147, siTMED2/10; n = 5794, siTMED2/10 + A939572; n = 4891, siTMED2/10 + Avasimibe from three replicates. The average Golgi fragments per cell are shown. ****p < 0.0001 (Unpaired two-tailed t-test). All scale bars are 10 µm.

**Figure S5.**
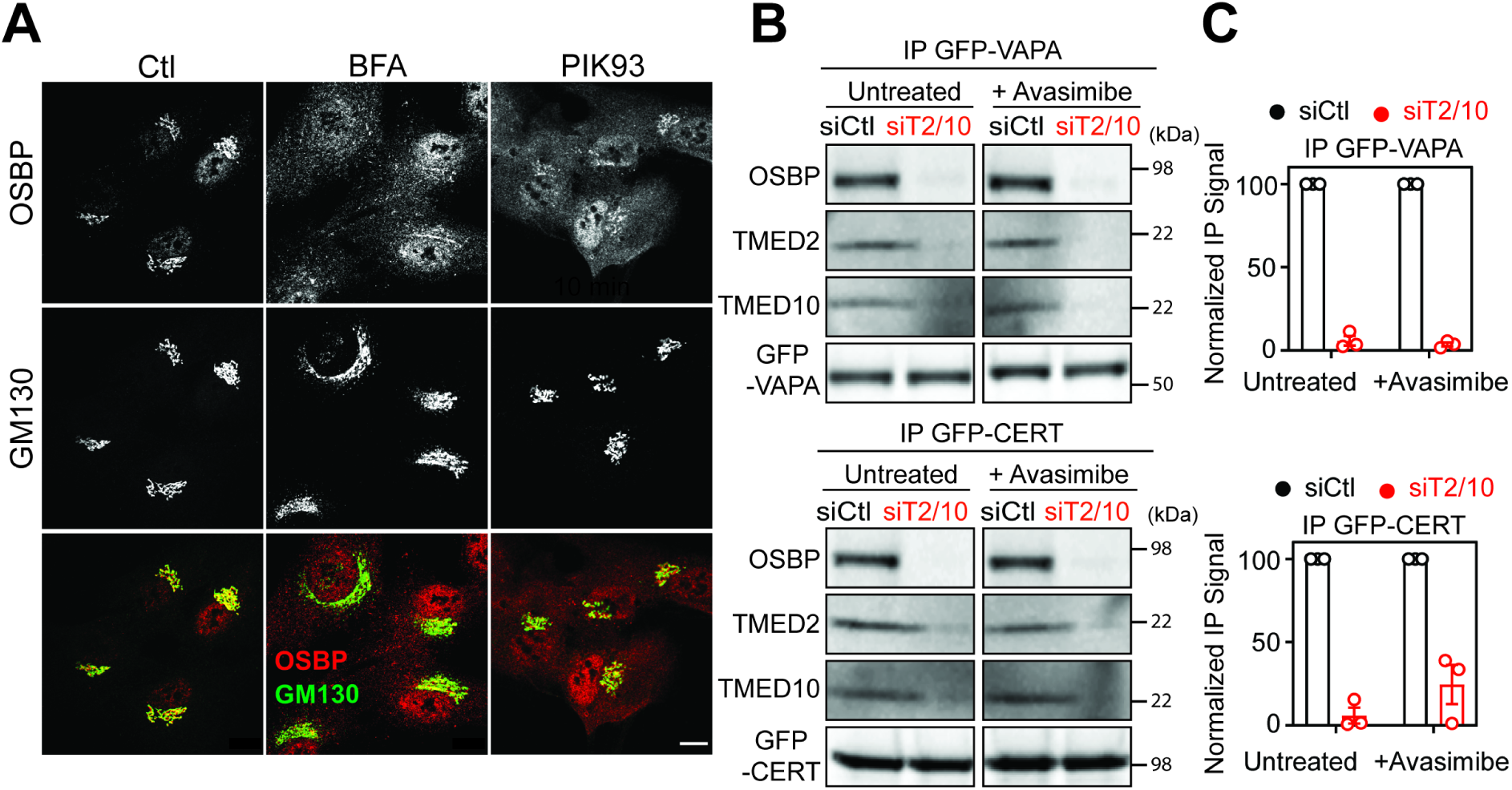
ACAT inhibition does not rescue ER-Golgi contacts in TMED2/10-silenced cells. **A)** Cells were treated with either BFA (5 µg/ml for 5 min) or PIK93 (4 µM for 10 min) before being fixed, permeabilized, and stained with anti-GM130 and anti-OSBP. Imaging was performed using laser scanning confocal microscopy. Scale bar is 10 µm. **B)** Western blot of GFP-VAPA and OSBP immunoprecipitated (IP) fractions. Before lysis and subsequent IPs, control and silenced cells were treated with Avasimibe (10 µM) for 24 h. **C)** The co-IP signal from the blot was set to 100 in siCtl cells. Results are mean ± SEM (n = 3).

## References

Abrami, L., Bischofberger, M., Kunz, B., Groux, R., & van der Goot, F. G. (2010). Endocytosis of the Anthrax Toxin Is Mediated by Clathrin, Actin and Unconventional Adaptors. PLoS Pathogens, 6(3), e1000792. https://doi.org/10.1371/journal.ppat.1000792

Abrami, L., Kunz, B., & Gisou van der Goot, F. (2010). Anthrax toxin triggers the activation of src-like kinases to mediate its own uptake. Proceedings of the National Academy of Sciences, 107(4), 1420–1424. https://doi.org/10.1073/pnas.0910782107

Abrami, L., Kunz, B., Iacovache, I., & van der Goot, F. G. (2008). Palmitoylation and ubiquitination regulate exit of the Wnt signaling protein LRP6 from the endoplasmic reticulum. Proceedings of the National Academy of Sciences, 105(14), 5384–5389. https://doi.org/10.1073/pnas.0710389105

Abrami, L., Leppla, S. H., & van der Goot, F. G. (2006). Receptor palmitoylation and ubiquitination regulate anthrax toxin endocytosis. Journal of Cell Biology, 172(2), 309–320. https://doi.org/10.1083/jcb.200507067

Abrami, L., Lindsay, M., Parton, R. G., Leppla, S. H., & van der Goot, F. G. (2004). Membrane insertion of anthrax protective antigen and cytoplasmic delivery of lethal factor occur at different stages of the endocytic pathway. Journal of Cell Biology, 166(5), 645–651. https://doi.org/10.1083/jcb.200312072

Abrami, L., Liu, S., Cosson, P., Leppla, S. H., & van der Goot, F. G. (2003). Anthrax toxin triggers endocytosis of its receptor via a lipid raft–mediated clathrin-dependent process. Journal of Cell Biology, 160(3), 321–328. https://doi.org/10.1083/jcb.200211018

Adarska, P., Wong-Dilworth, L., & Bottanelli, F. (2021). ARF GTPases and Their Ubiquitous Role in Intracellular Trafficking Beyond the Golgi. Frontiers in Cell and Developmental Biology, 9, 679046. https://doi.org/10.3389/fcell.2021.679046

Antonny, B., Bigay, J., & Mesmin, B. (2018). The Oxysterol-Binding Protein Cycle: Burning Off PI(4)P to Transport Cholesterol. Annual Review of Biochemistry, 87(1), 809–837. https://doi.org/10.1146/annurev-biochem-061516-044924

Bausch-Fluck, D., Hofmann, A., Bock, T., Frei, A. P., Cerciello, F., Jacobs, A., Moest, H., Omasits, U., Gundry, R. L., Yoon, C., Schiess, R., Schmidt, A., Mirkowska, P., Härtlová, A., Van Eyk, J. E., Bourquin, J.-P., Aebersold, R., Boheler, K. R., Zandstra, P., & Wollscheid, B. (2015). A Mass Spectrometric-Derived Cell Surface Protein Atlas. PLOS ONE, 10(4), e0121314. https://doi.org/10.1371/journal.pone.0121314

Bindea, G., Galon, J., & Mlecnik, B. (2013). CluePedia Cytoscape plugin: Pathway insights using integrated experimental and in silico data. Bioinformatics, 29(5), 661–663. https://doi.org/10.1093/bioinformatics/btt019

Bürgi, J., Abrami, L., Castanon, I., Abriata, L. A., Kunz, B., Yan, S. E., Lera, M., Unger, S., Superti-Furga, A., Peraro, M. D., Gaitan, M. G., & van der Goot, F. G. (2020). Ligand Binding to the Collagen VI Receptor Triggers a Talin-to-RhoA Switch that Regulates Receptor Endocytosis. Developmental Cell, 53(4), 418–430.e4. https://doi.org/10.1016/j.devcel.2020.04.015

Canals, D., Salamone, S., & Hannun, Y. A. (2018). Visualizing bioactive ceramides. Chemistry and Physics of Lipids, 216, 142–151. https://doi.org/10.1016/j.chemphyslip.2018.09.013

Capasso, S., Sticco, L., Rizzo, R., Pirozzi, M., Russo, D., Dathan, N. A., Campelo, F., Galen, J., Hölttä-Vuori, M., Turacchio, G., Hausser, A., Malhotra, V., Riezman, I., Riezman, H., Ikonen, E., Luberto, C., Parashuraman, S., Luini, A., & D’Angelo, G. (2017). Sphingolipid metabolic flow controls phosphoinositide turnover at the *trans* -Golgi network. The EMBO Journal, 36(12), 1736–1754. https://doi.org/10.15252/embj.201696048

Castillon, G. A., Aguilera-Romero, A., Manzano-Lopez, J., Epstein, S., Kajiwara, K., Funato, K., Watanabe, R., Riezman, H., & Muñiz, M. (2011). The yeast p24 complex regulates GPI-anchored protein transport and quality control by monitoring anchor remodeling. Molecular Biology of the Cell, 22(16), 2924–2936. https://doi.org/10.1091/mbc.e11-04-0294

Contreras, F.-X., Ernst, A. M., Haberkant, P., Björkholm, P., Lindahl, E., Gönen, B., Tischer, C., Elofsson, A., von Heijne, G., Thiele, C., Pepperkok, R., Wieland, F., & Brügger, B. (2012). Molecular recognition of a single sphingolipid species by a protein’s transmembrane domain. Nature, 481(7382), 525–529. https://doi.org/10.1038/nature10742

de la Mora, E., Dezi, M., Di Cicco, A., Bigay, J., Gautier, R., Manzi, J., Polidori, J., Castaño-Díez, D., Mesmin, B., Antonny, B., & Lévy, D. (2021). Nanoscale architecture of a VAP-A-OSBP tethering complex at membrane contact sites. Nature Communications, 12(1), 3459. https://doi.org/10.1038/s41467-021-23799-1

Donaldson, J. G., Honda, A., & Weigert, R. (2005). Multiple activities for Arf1 at the Golgi complex. Biochimica et Biophysica Acta (BBA) - Molecular Cell Research, 1744(3), 364–373. https://doi.org/10.1016/j.bbamcr.2005.03.001

Dong, R., Saheki, Y., Swarup, S., Lucast, L., Harper, J. W., & De Camilli, P. (2016). Endosome-ER Contacts Control Actin Nucleation and Retromer Function through VAP-Dependent Regulation of PI4P. Cell, 166(2), 408–423. https://doi.org/10.1016/j.cell.2016.06.037

Dorfer, V., Pichler, P., Stranzl, T., Stadlmann, J., Taus, T., Winkler, S., & Mechtler, K. (2014). MS Amanda, a Universal Identification Algorithm Optimized for High Accuracy Tandem Mass Spectra. Journal of Proteome Research, 13(8), 3679–3684. https://doi.org/10.1021/pr500202e

Emery, G., Gruenberg, J., & Rojo, M. (1999). The p24 family of transmembrane proteins at the interface between endoplasmic reticulum and Golgi apparatus. Protoplasma, 207(1–2), 24–30. https://doi.org/10.1007/BF01294710

Friebe, S., van der Goot, F., & Bürgi, J. (2016). The Ins and Outs of Anthrax Toxin. Toxins, 8(3), 69. https://doi.org/10.3390/toxins8030069

Gommel, D., Orci, L., Emig, E. M., Hannah, M. J., Ravazzola, M., Nickel, W., Helms, J. B., Wieland, F. T., & Sohn, K. (1999). P24 and p23, the major transmembrane proteins of COPI-coated transport vesicles, form hetero-oligomeric complexes and cycle between the organelles of the early secretory pathway. FEBS Letters, 447(2–3), 179–185. https://doi.org/10.1016/S0014-5793(99)00246-X

Gommel, D. U. (2001). Recruitment to Golgi membranes of ADP-ribosylation factor 1 is mediated by the cytoplasmic domain of p23. The EMBO Journal, 20(23), 6751–6760. https://doi.org/10.1093/emboj/20.23.6751

Hanada, K., Kumagai, K., Tomishige, N., & Yamaji, T. (2009). CERT-mediated trafficking of ceramide. Biochimica et Biophysica Acta (BBA) - Molecular and Cell Biology of Lipids, 1791(7), 684–691. https://doi.org/10.1016/j.bbalip.2009.01.006

Hanada, K., Kumagai, K., Yasuda, S., Miura, Y., Kawano, M., Fukasawa, M., & Nishijima, M. (2003). Molecular machinery for non-vesicular trafficking of ceramide. Nature, 426(6968), 803–809. https://doi.org/10.1038/nature02188

Hsu, N.-Y., Ilnytska, O., Belov, G., Santiana, M., Chen, Y.-H., Takvorian, P. M., Pau, C., van der Schaar, H., Kaushik-Basu, N., Balla, T., Cameron, C. E., Ehrenfeld, E., van Kuppeveld, F. J. M., & Altan-Bonnet, N. (2010). Viral Reorganization of the Secretory Pathway Generates Distinct Organelles for RNA Replication. Cell, 141(5), 799–811. https://doi.org/10.1016/j.cell.2010.03.050

Jiménez-Rojo, N., Leonetti, M. D., Zoni, V., Colom, A., Feng, S., Iyengar, N. R., Matile, S., Roux, A., Vanni, S., Weissman, J. S., & Riezman, H. (2020). Conserved Functions of Ether Lipids and Sphingolipids in the Early Secretory Pathway. Current Biology, 30(19), 3775–3787.e7. https://doi.org/10.1016/j.cub.2020.07.059

Kamentsky, L., Jones, T. R., Fraser, A., Bray, M.-A., Logan, D. J., Madden, K. L., Ljosa, V., Rueden, C., Eliceiri, K. W., & Carpenter, A. E. (2011). Improved structure, function and compatibility for CellProfiler: Modular high-throughput image analysis software. Bioinformatics, 27(8), 1179–1180. https://doi.org/10.1093/bioinformatics/btr095

Klimpel, K. R., Molloy, S. S., Thomas, G., & Leppla, S. H. (1992). Anthrax toxin protective antigen is activated by a cell surface protease with the sequence specificity and catalytic properties of furin. Proceedings of the National Academy of Sciences, 89(21), 10277–10281. https://doi.org/10.1073/pnas.89.21.10277

Kong, A. T., Leprevost, F. V., Avtonomov, D. M., Mellacheruvu, D., & Nesvizhskii, A. I. (2017). MSFragger: Ultrafast and comprehensive peptide identification in mass spectrometry–based proteomics. Nature Methods, 14(5), 513–520. https://doi.org/10.1038/nmeth.4256

Krantz, B. A., Finkelstein, A., & Collier, R. J. (2006). Protein Translocation through the Anthrax Toxin Transmembrane Pore is Driven by a Proton Gradient. Journal of Molecular Biology, 355(5), 968–979. https://doi.org/10.1016/j.jmb.2005.11.030

Larios, J., Mercier, V., Roux, A., & Gruenberg, J. (2020). ALIX- and ESCRT-III–dependent sorting of tetraspanins to exosomes. Journal of Cell Biology, 219(3). https://doi.org/10.1083/jcb.201904113

Levental, I., Levental, K. R., & Heberle, F. A. (2020). Lipid Rafts: Controversies Resolved, Mysteries Remain. Trends in Cell Biology, 30(5), 341–353. https://doi.org/10.1016/j.tcb.2020.01.009

Levine, T. P., & Munro, S. (2002). Targeting of Golgi-Specific Pleckstrin Homology Domains Involves Both PtdIns 4-Kinase-Dependent and -Independent Components. Current Biology, 12(9), 695–704. https://doi.org/10.1016/S0960-9822(02)00779-0

Liberali, P., Snijder, B., & Pelkmans, L. (2014). A Hierarchical Map of Regulatory Genetic Interactions in Membrane Trafficking. Cell, 157(6), 1473–1487. https://doi.org/10.1016/j.cell.2014.04.029

Lita, A., Pliss, A., Kuzmin, A., Yamasaki, T., Zhang, L., Dowdy, T., Burks, C., de Val, N., Celiku, O., Ruiz-Rodado, V., Nicoli, E.-R., Kruhlak, M., Andresson, T., Das, S., Yang, C., Schmitt, R., Herold-Mende, C., Gilbert, M. R., Prasad, P. N., & Larion, M. (2021). IDH1 mutations induce organelle defects via dysregulated phospholipids. Nature Communications, 12(1), 614. https://doi.org/10.1038/s41467-020-20752-6

Luo, J., Yang, H., & Song, B.-L. (2020). Mechanisms and regulation of cholesterol homeostasis. Nature Reviews Molecular Cell Biology, 21(4), 225–245. https://doi.org/10.1038/s41580-019-0190-7

Maekawa, M., & Fairn, G. D. (2015). Complementary probes reveal that phosphatidylserine is required for the proper transbilayer distribution of cholesterol. Journal of Cell Science, 128(7), 1422–1433. https://doi.org/10.1242/jcs.164715

Mañes, S., del Real, G., & Martínez-A, C. (2003). Pathogens: Raft hijackers. Nature Reviews Immunology, 3(7), 557–568. https://doi.org/10.1038/nri1129

Maxfield, F. R., & Wüstner, D. (2012). Analysis of Cholesterol Trafficking with Fluorescent Probes. In Methods in Cell Biology (Vol. 108, pp. 367–393). Elsevier. https://doi.org/10.1016/B978-0-12-386487-1.00017-1

Mesmin, B., Bigay, J., Moser von Filseck, J., Lacas-Gervais, S., Drin, G., & Antonny, B. (2013). A Four-Step Cycle Driven by PI(4)P Hydrolysis Directs Sterol/PI(4)P Exchange by the ER-Golgi Tether OSBP. Cell, 155(4), 830–843. https://doi.org/10.1016/j.cell.2013.09.056

Mesmin, B., Kovacs, D., & D’Angelo, G. (2019). Lipid exchange and signaling at ER–Golgi contact sites. Current Opinion in Cell Biology, 57, 8–15. https://doi.org/10.1016/j.ceb.2018.10.002

Mesquita, F. S., Goot, F. G., & Sergeeva, O. A. (2020). Mammalian membrane trafficking as seen through the lens of bacterial toxins. Cellular Microbiology, 22(4). https://doi.org/10.1111/cmi.13167

Milne, J. C., Furlong, D., Hanna, P. C., Wall, J. S., & Collier, R. J. (1994). Anthrax protective antigen forms oligomers during intoxication of mammalian cells. Journal of Biological Chemistry, 269(32), 20607–20612. https://doi.org/10.1016/S0021-9258(17)32036-7

Moreau, D., Vacca, F., Vossio, S., Scott, C., Colaco, A., Paz Montoya, J., Ferguson, C., Damme, M., Moniatte, M., Parton, R. G., Platt, F. M., & Gruenberg, J. (2019). Drug- induced increase in lysobisphosphatidic acid reduces the cholesterol overload in Niemann–Pick type C cells and mice. EMBO Reports, 20(7). https://doi.org/10.15252/embr.201847055

Nakatsu, F., & Kawasaki, A. (2021). Functions of Oxysterol-Binding Proteins at Membrane Contact Sites and Their Control by Phosphoinositide Metabolism. Frontiers in Cell and Developmental Biology, 9, 664788. https://doi.org/10.3389/fcell.2021.664788

Park, J. M., Greten, F. R., Li, Z.-W., & Karin, M. (2002). Macrophage Apoptosis by Anthrax Lethal Factor Through p38 MAP Kinase Inhibition. Science, 297(5589), 2048–2051. https://doi.org/10.1126/science.1073163

Ramachandran, R., Heuck, A. P., Tweten, R. K., & Johnson, A. E. (2002). Structural insights into the membrane-anchoring mechanism of a cholesterol-dependent cytolysin. Nature Structural Biology. https://doi.org/10.1038/nsb855

Rappsilber, J., Mann, M., & Ishihama, Y. (2007). Protocol for micro-purification, enrichment, pre-fractionation and storage of peptides for proteomics using StageTips. Nature Protocols, 2(8), 1896–1906. https://doi.org/10.1038/nprot.2007.261

Robinson, M. D., McCarthy, D. J., & Smyth, G. K. (2010). edgeR: A Bioconductor package for differential expression analysis of digital gene expression data. Bioinformatics, 26(1), 139–140. https://doi.org/10.1093/bioinformatics/btp616

Saito, S., Matsui, H., Kawano, M., Kumagai, K., Tomishige, N., Hanada, K., Echigo, S., Tamura, S., & Kobayashi, T. (2008). Protein Phosphatase 2Cɛ Is an Endoplasmic Reticulum Integral Membrane Protein That Dephosphorylates the Ceramide Transport Protein CERT to Enhance Its Association with Organelle Membranes. Journal of Biological Chemistry, 283(10), 6584–6593. https://doi.org/10.1074/jbc.M707691200

Schiavo, G., & van der Goot, F. G. (2001). The bacterial toxin toolkit. Nature Reviews Molecular Cell Biology, 2(7), 530–537. https://doi.org/10.1038/35080089

Schindelin, J., Arganda-Carreras, I., Frise, E., Kaynig, V., Longair, M., Pietzsch, T., Preibisch, S., Rueden, C., Saalfeld, S., Schmid, B., Tinevez, J.-Y., White, D. J., Hartenstein, V., Eliceiri, K., Tomancak, P., & Cardona, A. (2012). Fiji: An open-source platform for biological-image analysis. Nature Methods, 9(7), 676–682. https://doi.org/10.1038/nmeth.2019

Sergeeva, O. A., & van der Goot, F. G. (2019). Anthrax toxin requires ZDHHC5-mediated palmitoylation of its surface-processing host enzymes. Proceedings of the National Academy of Sciences, 116(4), 1279–1288. https://doi.org/10.1073/pnas.1812588116

Strating, J. R. P. M., & Martens, G. J. M. (2009). The p24 family and selective transport processes at the ER-Golgi interface. Biology of the Cell, 101(9), 495–509. https://doi.org/10.1042/BC20080233

Sun, J., & Jacquez, P. (2016). Roles of Anthrax Toxin Receptor 2 in Anthrax Toxin Membrane Insertion and Pore Formation. Toxins, 8(2), 34. https://doi.org/10.3390/toxins8020034

Whitfield, G. B., Brock, T. D., Ammann, A., Gottlieb, D., & Carter, H. E. (1955). Filipin, an Antifungal Antibiotic: Isolation and Properties. Journal of the American Chemical Society, 77(18), 4799–4801. https://doi.org/10.1021/ja01623a032

Zavodszky, E., & Hegde, R. S. (2019). Misfolded GPI-anchored proteins are escorted through the secretory pathway by ER-derived factors. ELife, 8, e46740. https://doi.org/10.7554/eLife.46740

Zheng, P., Gao, F., Deng, K., Gong, W., & Sun, Z. (2013). Expression, purification and preliminary X-ray crystallographic analysis of Arf1-GDP in complex with dimeric p23 peptide. Acta Crystallographica Section F Structural Biology and Crystallization Communications, 69(10), 1155–1158. https://doi.org/10.1107/S1744309113024330

